# Taxon-in-context reveals ecological individuality in the human gut microbiome

**DOI:** 10.64898/2026.10.22.760910

**Authors:** Joverlyn Gaudillo, Nicole Astrologo, Francesco Canonaco, Fabio Stella, Silvia Barbaresi, Teresa Zelante, Harm Wopereis, Yossi Cohen, Cyril Guyard, Jan Knol, Christophe Lay, Enzo Acerbi

## Abstract

Ecological theory holds that a microbe’s role depends on its surrounding community. Yet abundance profiles and fixed ecological groupings leave its individual context implicit. We introduce taxon-in-context, a reproducible representation of each microbe’s context within an individual’s microbiome. Across 7,700 metagenomes, we show that common gut species occupy different contexts in different people. This flexibility peaked in early life and declined through adulthood, a pattern reproduced in a larger genus-level compendium. For selected species, context within each person’s microbiome revealed age associations beyond species abundance and the person’s context mixture. In an independent longitudinal urinary tract infection cohort, microbes switched contexts within individuals. Taxon-in-context enabled recurrence prediction where an abundance based model performed near chance. Taxon-in-context and these findings expose a measurable, dynamic and individualized layer of microbiome organization, providing a basis for investigating why the same microbe or intervention may have different biological consequences across people.

## INTRODUCTION

The same gut microbe may participate in different biological processes in different individuals^1^. Its contribution to the host may therefore depend on the microbial assemblages in which it participates. The human gut microbiome is commonly described through the microbes it contains, their functions and their ecological groupings^2^. Taxonomic identity or abundance alone does not reveal a microbe’s ecological participation within a particular person’s microbiome. Resolving its context would allow us to examine how this participation varies between people and within a person over time.

Microbial guilds and enterosignatures have helped describe the ecological organization of microbiomes. Guilds group taxa by shared ecological or functional properties^3^, while enterosignatures describe each microbiome as a mixture of recurrent microbial assemblages^4^. Their usual representations apply the same guild membership or the same weight for a specific taxon within each enterosignature across samples and individuals. These shared memberships and weights describe population level organization. They do not ordinarily resolve the context in which each microbe participates within a particular individual’s microbiome.

The distinction resembles that between the possible meanings of a word and its meaning in a particular document. The word “cricket” can refer to an insect or a sport; the pattern of other words in the document indicates which meaning applies. Similarly, a microbe’s ecological context can be inferred from the microbial composition of the individual microbiome in which it occurs. Here we introduce taxon-in-context, a representation that makes this individual context explicit for each microbe. We define microbial contexts as recurrent taxonomic configurations inferred from microbiome composition across samples, providing a common reference for interpreting microbes within each person.

Latent Dirichlet Allocation^5^ (LDA) and related models have described microbiomes as mixtures of microbial assemblages^6,7^. However, instability across repeated fits is a longstanding reproducibility challenge for LDA: different assemblages can be recovered from the same data^8,9^. This makes it difficult to distinguish biological differences in a microbe’s context from differences introduced by the model. We developed Minutia Discovery (hereafter, Discovery), a robust and reproducible framework built on LDA that stabilizes microbial contexts and resolves the context of each taxon within each microbiome, enabling consistent comparisons across people and over time. Key terms are defined in Supplementary Table 1.

Applying Discovery to 7,700 human gut metagenomes from Curated Metagenomics Database^10^ (CMD) revealed widespread ecological context flexibility across the human gut microbiome. More than half of the species analysed were found to occupy different contexts across individuals, including many common gut microbes. This flexibility was greatest in early life and declined through adulthood, a pattern reproduced at genus level in a larger compendium.

Resolving these contexts made explicit a layer of age associated microbiome organization that remained implicit in abundance profiles. For selected species, the contexts they occupied distinguished samples from different host ages even after accounting for their abundance and the overall mixture of contexts in each sample. We also resolved how shared species were distributed across contexts in operationally defined typical and atypical microbiome configurations, connecting these broader aging associated patterns to the ecological settings of individual microbes. Context thus revealed how the ecological organization of individual microbes varies across the lifespan, beyond what is apparent from changes in their abundance alone.

Importantly, we detected context switching within individuals. In an independent longitudinal recurrent urinary tract infection (rUTI) cohort^11^, common gut microbes switched contexts within the same person during treatment and recovery. The microbial context of a species was therefore not fixed even within one host, raising the possibility that its biological contribution also changes as its ecological context changes.

Taxon-in-context also had clinical relevance. Within this cohort, taxon-in-context profiles predicted UTI recurrence better than matched continuous abundance profiles. Contextualizing microbes made clinically relevant organization more accessible from the same abundance data.

Taxon-in-context makes each microbe within each person’s microbiome an explicit unit of analysis. It provides a basis for assessing interventions through the ecological changes they induce. Prospective studies can test whether baseline context organization predicts intervention response. They can also test whether diet, biotics or other interventions shift the contexts occupied by resident or introduced microbes, even when their abundance changes little, and whether those shifts accompany differences in host response. This approach opens a route to understanding why the same microbe, or the same intervention, may have different biological consequences in different people

## RESULTS

### A reproducible context aware LDA framework identifies microbial contexts in the human gut

We applied the context aware LDA framework to 7,700 shotgun metagenomic profiles from the health-associated CMD reference cohort, spanning early life to later life. After taxonomic filtering and model selection, the reproducible solution identified four robust microbial contexts at species level from 275 retained taxa (Fig. 1c; Supplementary Fig. 1a). The full Discovery algorithm and model-selection procedure are described in <u>Supplementary Methods 1</u>.

**Fig. 1:**
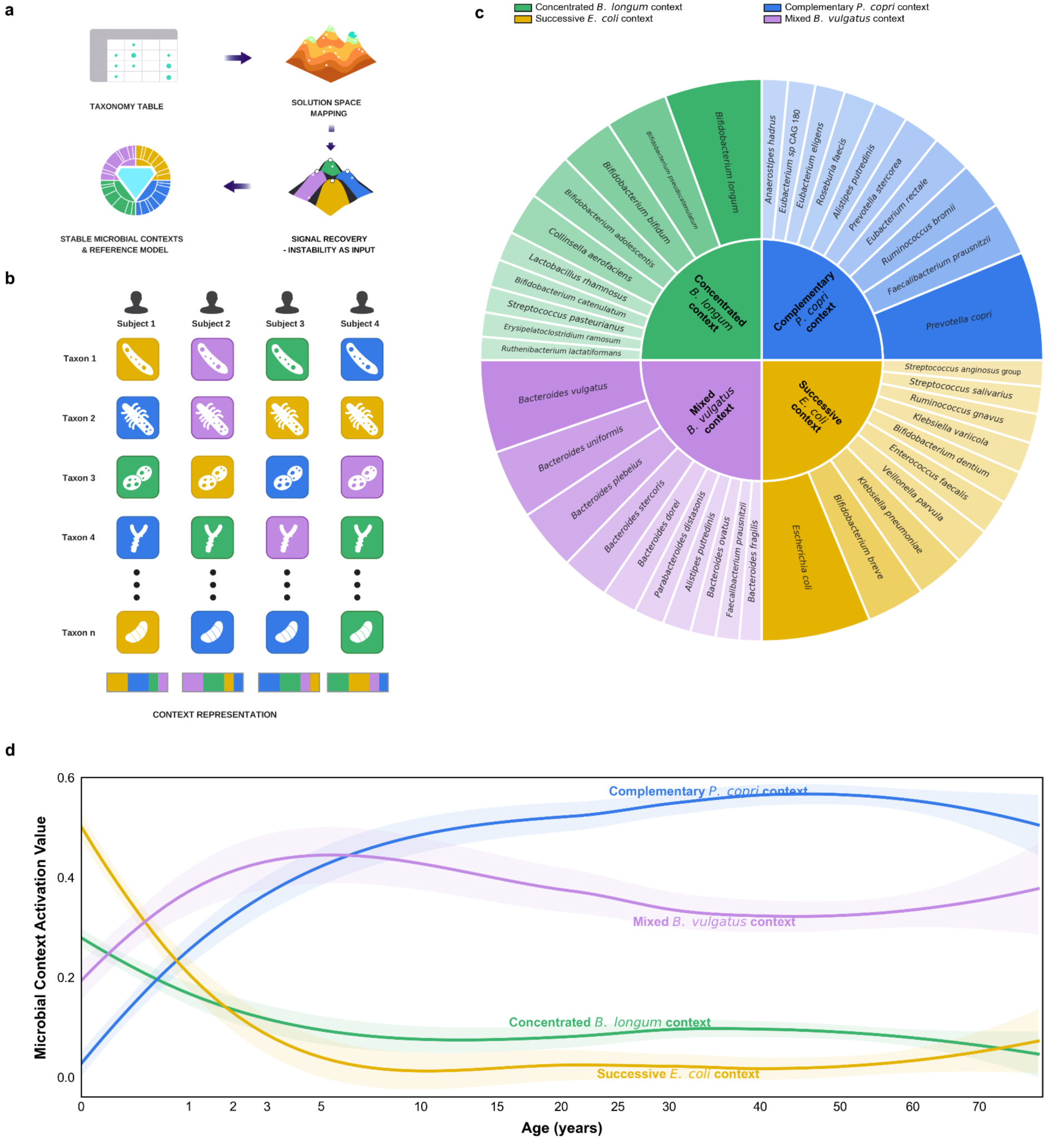
Discovery identifies species-level microbial contexts and age-associated activation patterns. **a,** Schematic of the Discovery workflow, in which independently initialized LDA fits are mapped in solution space to recover stable microbial contexts and a reference model. **b,** Schematic of sample-specific taxon-in-context assignment, showing the dominant inferred context for each retained taxon and the resulting sample-level context representation. **c,** Taxonomic composition of the four recovered microbial contexts. Inner sectors denote contexts and have equal angular width; outer sectors denote displayed taxa with non-zero context-composition weights. Outer-sector width and shade intensity represents within-context composition weights. **d,** Age-associated context activation profiles among 7,700 age-annotated CMD samples with matched context profiles. Context activations were normalized within each sample to sum to 1, averaged in consecutive age-ordered windows, and fitted using locally weighted linear regression (LOESS) on a signed-square-root age scale; shaded bands indicate pointwise 95% confidence intervals.

To facilitate interpretation, each context name combined its highest probability taxon with a descriptor of how its highest routed-mass contributor taxa (<u>Supplementary Methods 2</u>) were organized across microbiomes and life stages: Mixed *Bacteroides vulgatus*, Concentrated *Bifidobacterium longum*, Successive *Escherichia coli* and Complementary *Prevotella copri* (<u>Supplementary Methods 2</u>; Supplementary Figure 2; <u>Supplementary Results 1</u>). Complementary *P. copri* showed the strongest tendency for taxa to occupy the context together within the same samples. In Successive *E. coli*, different taxa contributed most at different life stages. Concentrated *B. longum* retained the same small set of leading contributors across life stages. Mixed *B. vulgatus* combined several patterns among taxon pairs

Independent repetitions of the complete Discovery workflow recovered the same four microbial context identities (Supplementary Fig. 3a,b; Supplementary Table 2). Among sample–taxon observations with valid assignments in all six Discovery runs, 98% received the same context assignment in every run (<u>Supplementary Methods 3</u>; <u>Supplementary Results 2</u>; Supplementary Fig. 3c,d; Supplementary Table 3).

We next repeated the Discovery analysis at the genus level in a larger compendium combining the Gut Microbiome Reference^4^ and Human Microbiome Compendium^12^ cohorts, referred to as GMR-Compendium (<u>Supplementary Methods 4</u>). Discovery recovered broadly comparable microbial contexts and age-associated patterns in this combined dataset (<u>Supplementary Results</u> <u>3</u>; Supplementary Fig. 1b; Supplementary Fig. 4a). We also compared contexts with enterosignatures^4^. Most microbial contexts are aligned with a specific enterosignature at the membership level (<u>Supplementary Results 4</u>). The two frameworks are closely related in construction, what differs is the unit of analysis: enterosignatures quantify which population-level components are active in a sample, whereas Discovery additionally represents, for each sample, the context that each individual taxon occupies.

### Microbial context activation describes lifespan microbiome development

Context activation profiles describe each microbiome as a mixture of microbial contexts, with probability weights indicating their relative contributions. We next examined whether these profiles captured known developmental structure in the gut microbiome (Fig. 1d).

Across age annotated samples, the Successive *E. coli* context was most prominent in early life and declined rapidly with age. The Concentrated *B. longum* context increased during infancy, while the Mixed *B. vulgatus* context became prominent after the first year of life and remained broadly active across later life stages. The Complementary *P. copri* context increased more gradually after childhood and became a major contributor to adult microbiome organization. These trajectories were reproduced at genus level in the GMR-Compendium (Supplementary Fig. 4b).

### Common gut microbes occupy different ecological contexts across individuals

We asked whether microbes remained anchored to a single context across individuals or occupied alternative contexts in different microbiomes. Across the 275 taxa retained in the CMD analysis, 126 remained associated with one context, whereas 149 (54%) were multicontext taxa. Their median prevalence across samples was higher than that of single-context taxa, 21.3% versus 10.4% (two-sided Mann-Whitney U test, *P* = 2. 3 × 10^-5^), while their mean relative abundance was comparable, 0.362% versus 0.352% (*P* = 0. 784). Several common gut commensals, including *Faecalibacterium prausnitzii*, *Blautia wexlerae*, *Bacteroides vulgatus* and *Eubacterium rectale*, were prevalent multicontext taxa.

### Ecological context flexibility peaks in early life and declines through adulthood

We next quantified whether ecological context flexibility followed reproducible lifespan structure. Low context entropy indicates that a taxon is consistently assigned to the same context across individuals of the same age, whereas high context entropy indicates that the taxon is distributed across multiple microbial contexts. In the study and subject controlled analysis, ecological context flexibility was highest in early life, at ages spanning the weaning window, and then declined to a substantially lower level in adulthood, where it continued to decrease steadily with age. Both the early life maximum and the adult decline were robust to the participant subsampling depth and to leaving out any single contributing study. The GMR-Compendium similarly showed elevated flexibility in early childhood and lower levels in adulthood (Supplementary Fig. 5). Directed transition networks (Fig. 2a) showed the densest and most bidirectional taxa-context reallocation in early childhood, and countervailing routes that reorganize taxa in later stages with little net change in sample-level context activation.

**Fig. 2:**
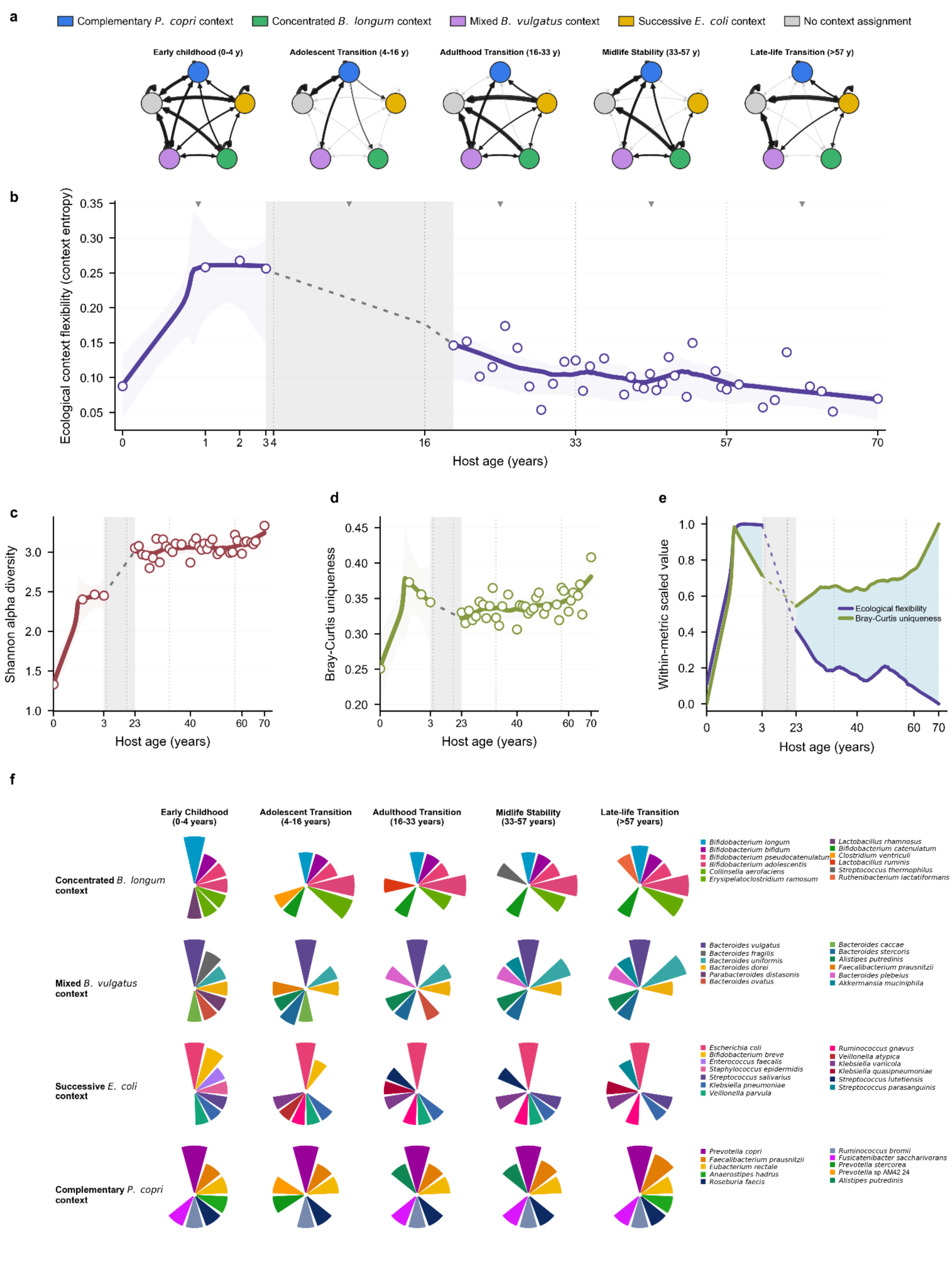
Age-associated patterns in microbial-context routing, ecological flexibility, diversity, uniqueness and taxon contributions. **a,** Directed microbial-context route networks across life stages. Nodes represent the microbial contexts and the no-context-assignment state; arrows connect context states observed for taxa across consecutive ages within each life stage. Black arrows denote routes that increased or emerged relative to the preceding life stage, whereas grey arrows denote routes that decreased or remained unchanged; arrow width reflects route strength. These cross-sectional networks summarize age-associated shifts in taxon context occupancy. **b,** Prevalence-weighted ecological context flexibility across age. Open circles show age-specific study-balanced estimates, solid lines show fitted trajectories and shaded bands indicate pointwise 95% bootstrap confidence intervals. The grey region denotes the lower-support interval. **c,d,** Shannon alpha diversity (**c**) and nearest-neighbour Bray-Curtis uniqueness (**d**) across age, using the same conventions as in **b**. **e,** Independently scaled ecological flexibility and Bray-Curtis uniqueness trajectories; blue shading shows the separation between curves. **f,** Taxon contributions to each microbial context across life stages. Petals show the seven taxa with the highest routed mass within each context–life-stage combination; petal length is scaled within each plot and colours identify taxa. Routed-mass calculation is described in <u>Supplementary Methods 2</u>.

The reduction in ecological flexibility (Fig. 2b) occurred alongside an overall increase in within-sample taxonomic diversity (Fig. 2c) and a distinct trajectory of nearest-neighbour Bray-Curtis uniqueness^13^ (Fig. 2d). When scaled within metric, flexibility and uniqueness were broadly concordant during early childhood but diverged after the adolescent interval (Fig. 2e). Flexibility progressively decreased, whereas uniqueness increased across adulthood and late life.

Context-specific routed mass share profiles, representing the fraction of abundance assigned to a context contributed by each taxon, showed that these lifespan changes were accompanied by redistribution of the microbes contributing to each context (Fig. 2f).

### Taxon-specific context occupancy improves age modelling

To determine whether the ecological context occupied by a taxon carried age-related information beyond taxon abundance, we performed taxon specific nested model comparisons (Supplementary Methods 5).

Among the 149 multicontext species, 93 were retained after requiring at least twenty samples in each occupied context, and 64 of these showed a significant improvement in model fit when taxon-in-context assignment was added to focal taxon abundance (*q* < 0. 01; mean abundance-controlled Δ*R* = 0. 062). These species are hereafter referred to as context-responsive taxa (<u>Supplementary Data 1)</u>.

A composition-controlled analysis then tested, across all 93 eligible taxa, whether context assignments were associated with age by accounting for both focal taxon abundance and the sample-level context activation profile. We assessed whether these associations were stronger than those obtained from random assignments generated according to each sample’s overall mixture of contexts. Forty-two taxa showed a significant empirical association after multiple-testing correction (*q_emp_* < 0. 05) together with a positive composition-controlled Δ*R*^2^.

We refer to these taxa as context pivots (<u>Supplementary Data 1</u>). The context pivots are dominated by fibre fermenting and cross feeding anaerobes including *Faecalibacterium prausnitzii*, *Ruminococcus torques*, *Anaerostipes hadrus*, and *Eubacterium eligens*. Thirty-seven taxa qualified as both context-responsive taxa and context pivots, whereas most broad-only taxa showed attenuation after composition adjustment, indicating that their assignment signal was largely shared with global context structure (<u>Supplementary Results 5</u>).

Across all samples, the 64 context-responsive taxa collectively accounted for an average of 41.3% of total relative abundance. Mean context entropy and routed mass share are compared in Fig. 3a, which separates flexible but low-contributing taxa from taxa that combine high context flexibility with substantial ecological weight.

**Fig. 3:**
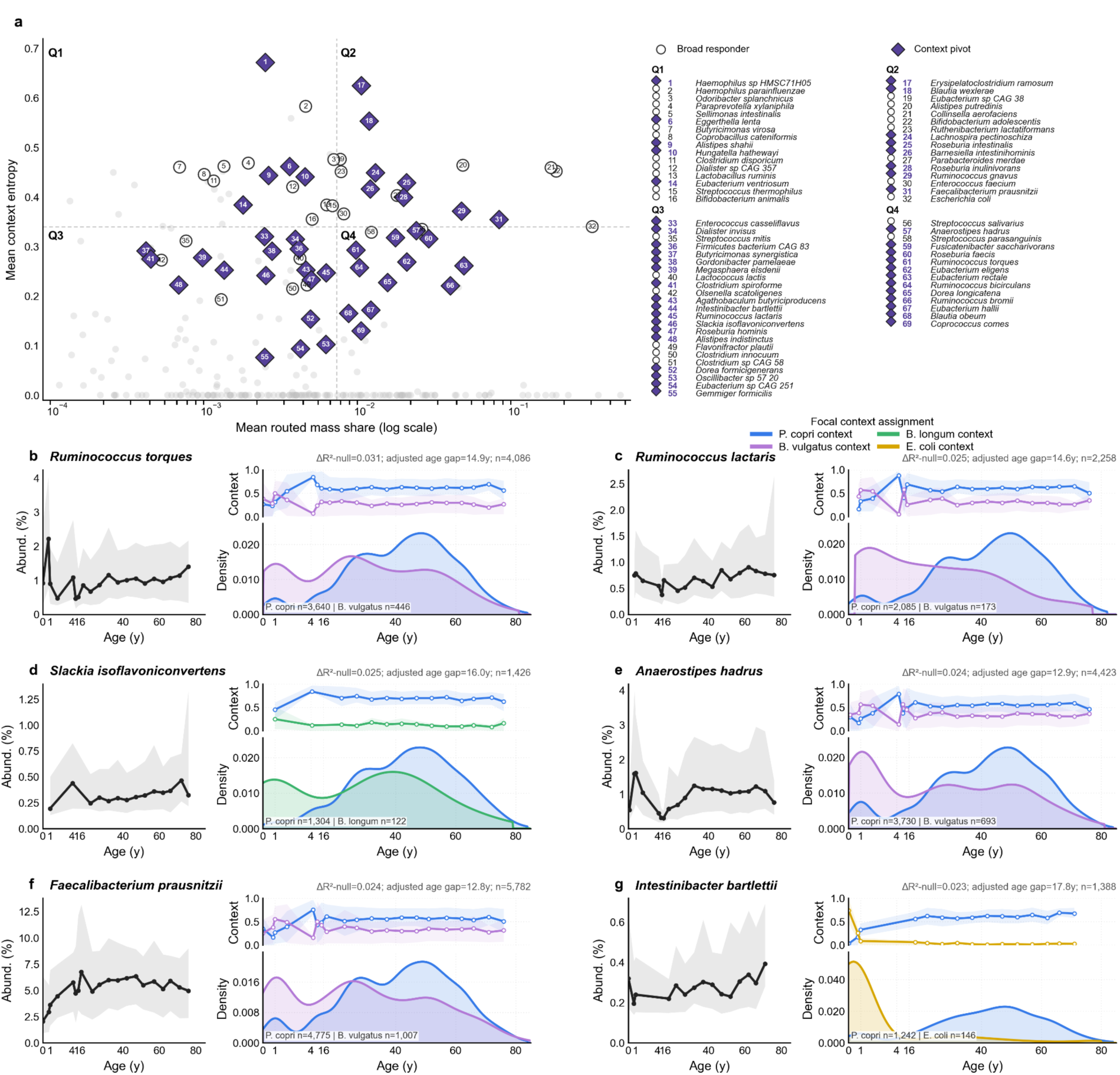
Taxon-in-context occupancy reveals age-associated structure within individual taxa. **a,** Mean context entropy versus mean routed-mass share across evaluated taxa. Open circles denote context-responsive taxa, purple diamonds denote context pivots and grey points denote other evaluated taxa**. b-g,** Age-associated profiles of six representative context pivots: **b,** *Ruminococcus torques*; **c,** *Ruminococcus lactaris*; **d,** *Slackia isoflavoniconvertens*; **e,** *Anaerostipes hadrus*; **f,** *Faecalibacterium prausnitzii*; and **g,** *Intestinibacter bartlettii*. For each taxon, the left plot shows median relative abundance with interquartile range, the upper-right plot shows mean sample context-composition weights with interquartile range, and the lower-right plot shows host-age density distributions by taxon-in-context assignment. ΔR²-null quantifies assignment-associated age information beyond focal abundance and sample context composition relative to the permutation null; adjusted age gap denotes the corresponding maximum model-adjusted between-context age difference. Age axes are display-transformed but labelled in years. Values of *n* denote retained sample-taxon observations.

Representative context pivots included *Ruminococcus torques*, *Ruminococcus lactaris*, *Slackia isoflavoniconvertens*, *Anaerostipes hadrus*, *Faecalibacterium prausnitzii* and *Intestinibacter bartlettii* (Fig. 3b-g). For example, *R. torques* and *F. prausnitzii* were assigned predominantly to the Complementary *P. copri* and Mixed *B. vulgatus* contexts, with adjusted between-context age gaps of approximately 14.9 and 12.8 years, respectively.

### Taxon-in-context occupancy is associated with delivery mode and microbiome configuration

We analyzed infants aged 0 to 1 year to test whether taxon-in-context assignment was associated with delivery mode (n = 1,650; 932 born by vaginal delivery [VD] and 718 by caesarean section [CS]) after controlling for species abundance. This identified 22 species with delivery mode associated context assignment after multiple testing correction (*q* < 0. 01; Fig. 4a).

**Fig. 4:**
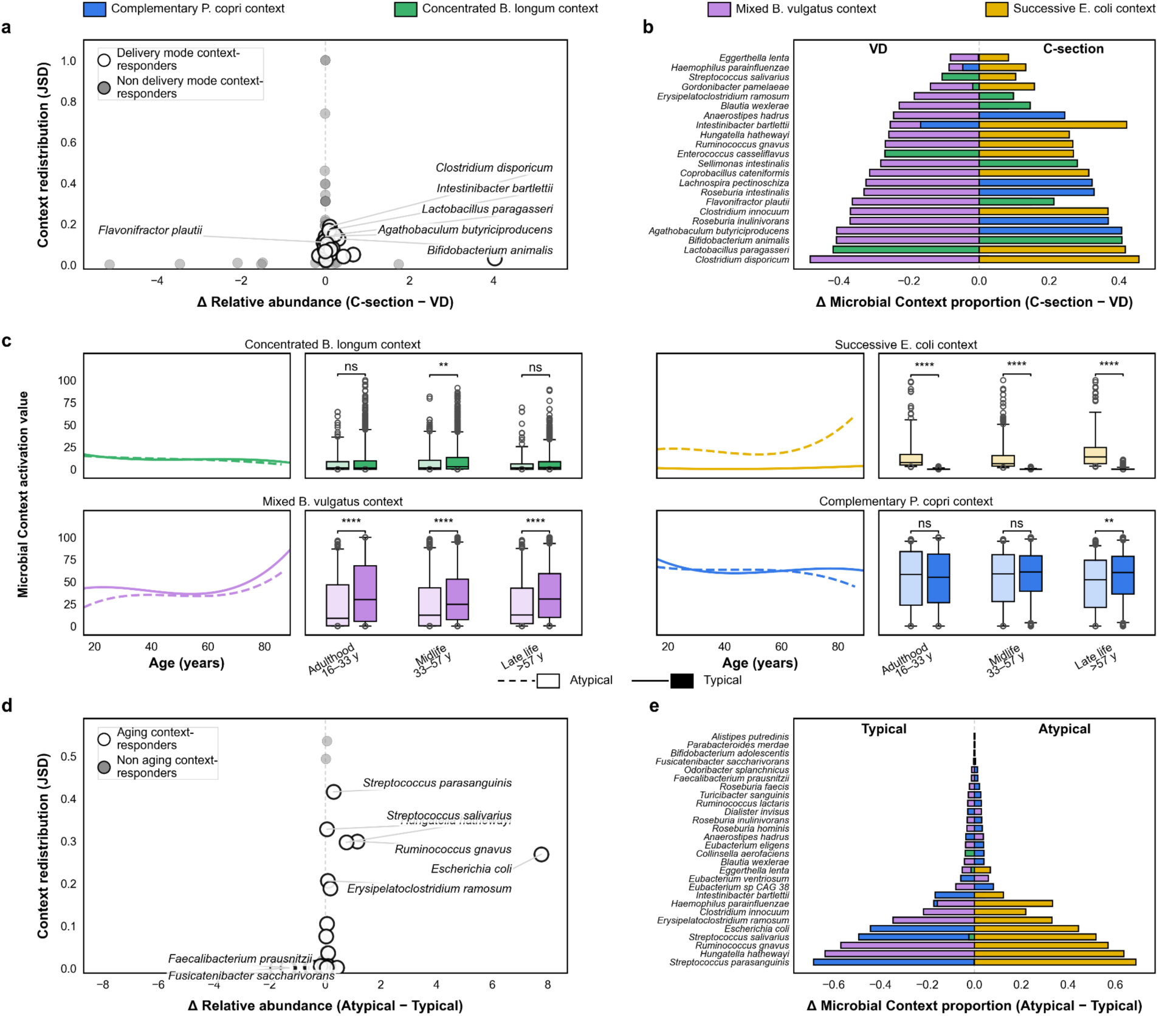
Delivery mode- and microbiome configuration-associated redistribution of taxa across microbial contexts. a,d, Differences in mean relative abundance (percentage points) versus context redistribution (base-2 Jensen–Shannon divergence) for delivery mode (C-section - VD; n = 1,650 samples, 932 VD and 718 C-section) and atypical versus typical configurations (atypical - typical; n = 744 and 4,278, respectively). Open circles highlight selected taxa; labelled taxa are selected examples. **b,e,** Context-specific assignment-proportion differences for the 22 and 28 highlighted taxa in the delivery-mode and aging-configuration comparisons, respectively; positive values indicate higher assignment proportions in C-section or atypical samples and negative values in VD or typical samples. **c,** Age-associated context activation in the same 5,022 samples aged 16-89 years. Solid and dashed lines show typical and atypical age patterns, respectively, using cubic fits to rolling maxima of age-specific mean activation. Boxes show medians and interquartile ranges; whiskers indicate the 5th-95th percentiles, with observations beyond the whiskers plotted individually. Comparisons use two-sided Mann–Whitney U tests with unadjusted *P* values: \**P* ≤ 0.05, \*\**P* ≤ 0.01, \*\*\**P* ≤ 0.001, \*\*\*\**P* ≤ 0.0001; ns, *P* > 0.05.

Taxa in CS infants were more frequently assigned to the Successive *E. coli* context, whereas taxa in VD infants were more frequently assigned to the Mixed *B. vulgatus* context (Fig. 4b). Several delivery mode associated species showed marked context redistribution despite relatively small differences in abundance. Thus, delivery mode was reflected not only in taxon abundance, but also in the microbial contexts to which taxa were assigned.

We next asked whether taxon-in-context assignment also distinguished aging-associated microbiome configurations. The atypical configuration was operationally defined by elevated activation of the Successive *E. coli* context, as defined in the Methods, outside early life, where this context is less expected and is enriched in facultative anaerobic/aerobic and opportunistic taxa^14^ (Fig. 4c). This classification was applied from adulthood transition through late life, yielding 4,278 typical samples and 744 atypical samples.

After adjusting for species abundance and applying multiple testing corrections, 28 species showed context redistribution associated with typical versus atypical configurations (*q* < 0. 05; Fig. 4d). These species were more frequently assigned to the Successive *E. coli* context in samples classified as atypical (Fig. 4e).

### Gut microbes switch contexts within individuals during UTI treatment and recovery

To determine whether multicontext organization observed across individuals also occurs within the same host over time, we analyzed an independent longitudinal recurrent urinary tract infection (rUTI) cohort^11^. The CMD-learned contexts were projected onto 480 stool metagenomes from 106 participants after taxonomic harmonization; the 105 participants with at least two samples were retained.

Within-person switching was widespread: 90 of 105 participants (85.7%; 95% CI, 79.1 to 92.4%) showed at least one taxon that occupied different microbial contexts across their longitudinal samples, with a median of five switching taxa per participant (interquartile range, 2-9; Fig. 5a). Within participants, switching microbes were usually restricted to two contexts, 83.2% of context-switching events followed the Mixed *B. vulgatus* - Complementary *P. copri* axis, with transitions occurring in both directions (Fig. 5b).

**Fig. 5:**
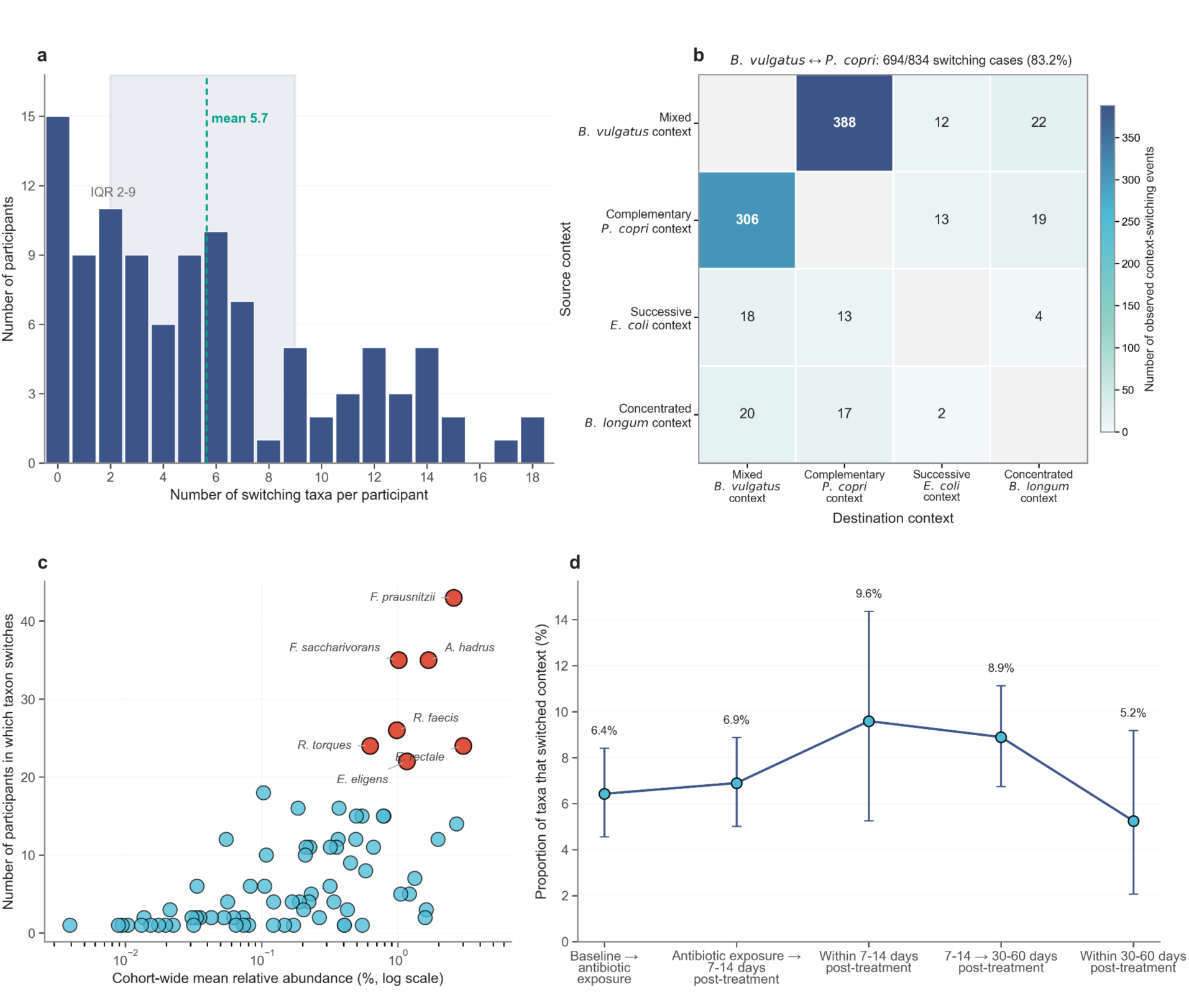
Gut microbes switch ecological contexts within individuals across UTI treatment and recovery. **a,** Distribution of within-participant context switching across the full longitudinal series. The dashed line shows the cohort mean (5.7 taxa) and shading the interquartile range (2-9 taxa). **b,** Directional strict context-switching events between consecutive available samples. **c,** Cohort-wide mean relative abundance versus the number of participants in which each taxon switched context. **d,** Proportion of participant–taxon transitions classified as strict context switches across selected treatment-related sampling intervals. Clinical periods were defined as baseline (T1), antibiotic exposure (T2-T3), 7-14 days post-treatment (T4-T5), and 30-60 days post-treatment (T6-T7). Strict context switching was defined as a change in the assigned microbial context of the same taxon between consecutive available samples from the same participant, requiring a valid context assignment at both samples (see Methods for the full definition). Switching proportions were calculated as the number of strict context-switching events divided by the number of participant-taxon transitions with valid context assignments at both consecutive samples within each displayed interval. Points show observed proportions and error bars indicate patient-cluster bootstrap 95% CI based on 2,000 replicates.

The most recurrent switching taxa were *Faecalibacterium prausnitzii* (43 participants; cohort-wide mean relative abundance, 2.56%), *Fusicatenibacter saccharivorans* (35; 1.01%), *Anaerostipes hadrus* (35; 1.67%), *Roseburia faecis* (26; 0.98%), *Ruminococcus torques* (24; 0.62%), *Eubacterium rectale* (24; 3.01%) and *Eubacterium eligens* (22; 1.16%). Several frequent switchers were quantitatively prominent members of the gut community (Fig. 5c). Across the selected clinical periods, strict context-switching rates were 6.4% from baseline to antibiotic exposure, 6.9% from antibiotic exposure to 7–14 days post-treatment, 9.6% within 7–14 days post-treatment, 8.9% from 7–14 to 30–60 days post-treatment and 5.2% within 30–60 days post-treatment (Fig. 5d).

### CMD-derived taxon-in-context profiles predict recurrent UTI outcome in an independent longitudinal cohort

We next tested whether taxon-in-context profiles could predict recurrence in the independent rUTI cohort. These profiles used the contexts previously learned from CMD, requiring no disease specific context discovery. The analysis was restricted to each participant’s first two observed timepoints to limit outcome related follow up information. This retained 105 participants and 210 metagenomes. Retained samples were concentrated at enrollment and end of treatment, and timing-only models separated outcomes substantially less well than the context model, although practical equivalence to chance was not established (<u>Supplementary Methods 6</u>; <u>Supplementary Results 6</u>; Supplementary Table 4). All representations in the primary recurrence analysis used the 223 taxa represented in both the CMD reference and the episode-1 modeling dataset.

Taxon-in-context assignment discriminated recurrence with a cross validated area under the receiver operating characteristic curve (AUROC) of 0.707 (95% CI, 0.600 to 0.805), compared with 0.543 (95% CI, 0.424 to 0.663) for the matched abundance model (Fig. 6a; Supplementary Table 5). Context exceeded centred log-ratio (CLR) abundance by 0.164 AUROC units (95% CI, 0.053 to 0.281).

**Fig. 6:**
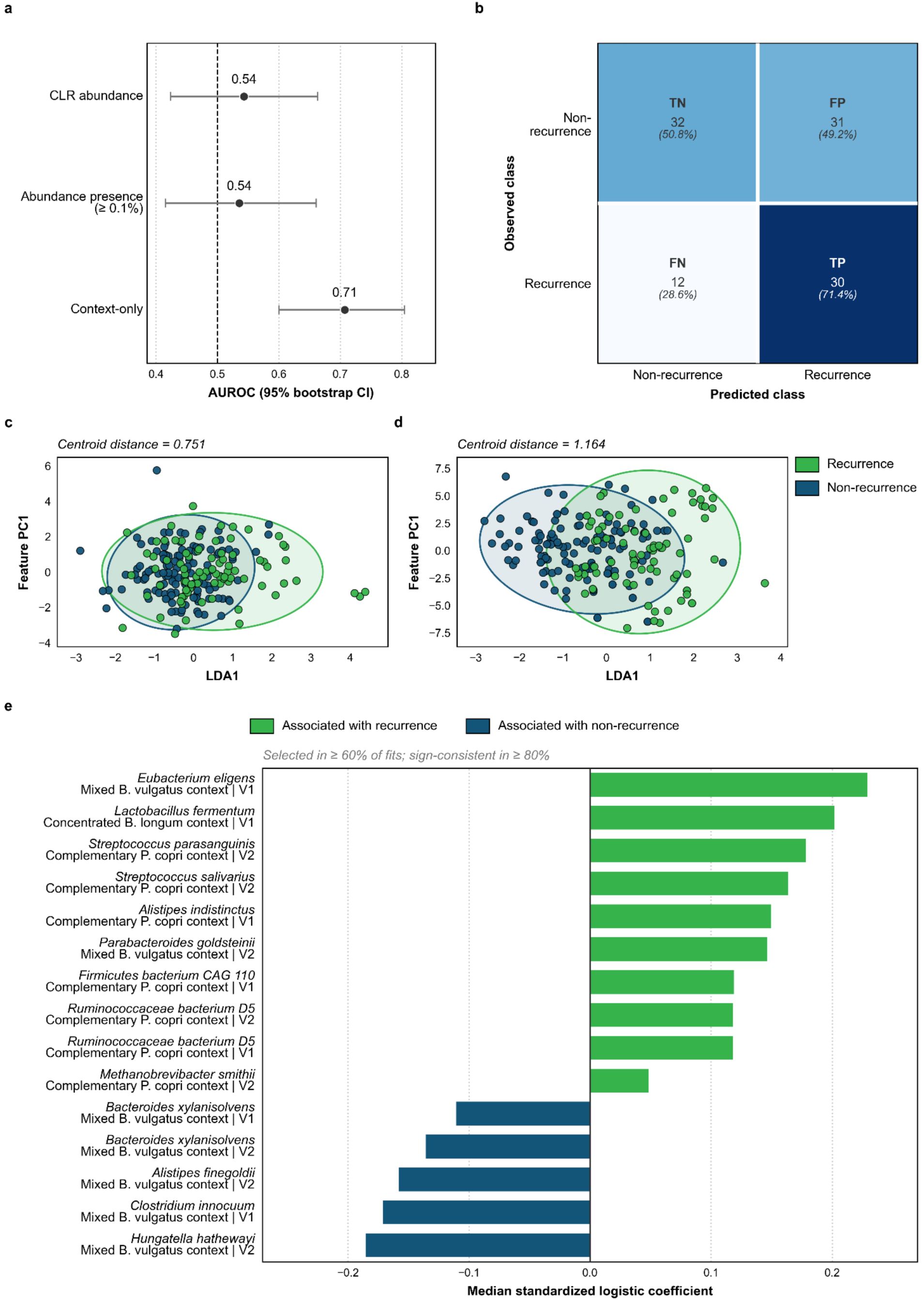
Prediction of recurrent urinary tract infection from abundance and microbial-context features at ordinal visits 1–2. **a,** Participant-level out-of-fold AUROC for matched CLR abundance, thresholded presence (≥0.1%) and context-only models in 105 participants. Error bars show conditional 95% confidence intervals from 2,000 participant-level bootstrap resamples of pooled outer-fold probabilities without refitting; the dashed line indicates chance discrimination (0.5). **b,** Context-only confusion matrix using outer-test predictions and fold-specific thresholds selected from inner out-of-fold training predictions. **c,d,** Descriptive sample-level projections of raw abundance (**c**) and one-hot-encoded context states, including a no-assignment state (**d**), for 24 context-represented taxa selected by total abundance across input samples (209 samples; 105 participants). Features were standardized before projection. Axes show supervised linear discriminant axis 1 (LDA1), fitted using recurrence labels, and the first principal component of standardized features (Feature PC1). Points represent samples; ellipses summarize group covariance. Centroid distances are Euclidean distances between group means after scaling each projected axis to unit standard deviation. Separation is descriptive. **e,** Stable context-only predictors, selected in ≥60% of outer fits with ≥80% sign consistency among nonzero coefficients. Bars show median standardized logistic-regression coefficients; positive and negative values indicate recurrence and non-recurrence associations, respectively. Additional comparator and stable-feature results are reported in <u>Supplementary Results 6</u> and Supplementary Tables 5 and 7.

The context-only model correctly identified 30 of 42 recurrence cases and 32 of 63 non-recurrence cases (Fig. 6b). Sensitivity was 0.714 (95% CI, 0.571 to 0.853), whereas specificity was 0.508 (95% CI, 0.387 to 0.632). The corresponding positive predictive value was 0.492, negative predictive value was 0.727, and balanced accuracy was 0.611.

In terms of descriptive comparisons, supervised sample-level projections of 24 selected taxa showed clearer separation between samples from participants with and without recurrence using taxon-in-context profiles than abundance profiles (Fig. 6c-d; <u>Supplementary Results 6</u>). Because context assignment requires taxa to meet the 0.1% abundance floor, we also evaluated a threshold-matched presence representation. This model achieved an AUROC of 0.536 (95% CI, 0.415 to 0.661), with an estimated context-versus-presence difference of 0.172 (95% CI, 0.060 to 0.287; Supplementary Table 5). In a separate sensitivity analysis using a common 226-taxon universe, taxon-in-context achieved higher AUROC than both CLR abundance and thresholded presence across all five linear classifiers when participant-level out-of-fold scores were averaged across 20 repetitions of nested cross-validation (<u>Supplementary Results 6</u>; Supplementary Table 6). Fifteen taxon-in-context-visit predictive features met a pre-specified stability criterion (Fig. 6e; <u>Supplementary Results 6</u>; Supplementary Table 7).

Together, these findings support taxon-in-context as an interpretable representation for recurrence prediction. It improved discrimination relative to CLR abundance in the primary analysis, with a consistent advantage across classifiers in sensitivity analyses.

## DISCUSSION

Microbial guilds and enterosignatures describe recurrent organization through shared memberships or taxonomic weights^3,4^. Taxon-in-context makes the ecological setting of each microbe within each person’s microbiome explicit. A person’s mixture of contexts describes the microbiome as a whole; the context occupied by a particular taxon locates that microbe within it. Our findings reveal ecological individuality: the same gut microbes can occupy different ecological contexts in different people. Some microbes also exhibited context switching within individuals over time. Discovery provides the reproducible reference needed to compare these settings, making ecological individuality accessible from the same abundance data in which it was previously implicit.

*Bifidobacterium adolescentis* illustrates the biological relevance of resolving a species within its community. It occupied either the Concentrated *B. longum* or the Complementary *P. copri* context. Given its capacity to utilize plant carbohydrates and support butyrate production through cross feeding^15^, these alternative settings suggest that its contribution may depend on the substrates and microbial partners available within each person’s microbiome. The assignments identify where such interactions can be investigated, connecting a species’ known capabilities to the community in which they might be expressed. Further biological interpretations are provided in <u>Supplementary Discussions</u>.

Context flexibility followed a reproducible pattern across the lifespan. It was greatest in early life and declined through adulthood, with the pattern reproduced at genus level in a larger compendium. The early life maximum is consistent with rapid community assembly as milk feeding gives way to a broader diet^16^. Delivery mode associated redistribution further suggests that early colonization is reflected in the contexts occupied by microbes. In adults, declining flexibility accompanied increasing compositional uniqueness^13^ and sustained alpha diversity. Thus, microbiomes could differ increasingly in composition while particular shared species occupied more consistent contexts across people of similar age. An increasingly individual microbiome can therefore retain common ecological settings for its resident microbes.

The taxon specific age associations further show why a microbe’s ecological position matters. Context-responsive taxa can index the broader microbiome configuration. For context pivots, including *Faecalibacterium prausnitzii*, people of different ages differed in the ecological settings occupied by these species. These associations remained after accounting for species abundance and the modelled effects of each person’s overall context mixture. Whether context pivots initiate or maintain a context requires experimental testing. In the operationally defined atypical configuration, multiple shared species were more often assigned to the Successive *E. coli* context, showing that the contrast extended beyond *E. coli* abundance alone.

The longitudinal rUTI cohort^11^ extends this perspective from differences between people to changes within an individual. Common commensals, including *F. prausnitzii*, switched contexts during treatment and recovery. A species’ ecological setting can therefore vary within the same host, raising the question of whether its biological contribution changes with it. Denser longitudinal observations could establish whether these changes are transient or persist after recovery. Switching counts, directions and specific transitions did not predict recurrence in this cohort.

In the same rUTI cohort, taxon-in-context configurations from each participant’s first two observed timepoints predicted recurrence under internal cross validation. Contexts learned in CMD transferred without refitting to this independent cohort. Taxon-in-context enabled recurrence discrimination where the matched continuous abundance model performed near chance. Its advantage over taxon presence was consistent across multiple classifiers.

Particular combinations of microbe and context were associated with recurrence, whereas others were associated with non recurrence. Prospective validation in independent cohorts with predefined collection times remains necessary.

Several limitations guide the biological interpretation. The lifespan analyses describe variation across people and do not establish how an individual’s microbiome ages. Study balanced resampling supports the aggregate flexibility curve, but taxon specific age and delivery mode associations may still reflect study composition or incompletely measured host, dietary and geographic factors. Likewise, typical and atypical describe context defined configurations, not independently validated health states. Sample size support criteria restricted the primary flexibility curve and taxon specific age comparisons to adequately represented groups. Larger, more evenly sampled cohorts, denser longitudinal series and explicit assignment probabilities would help distinguish rarer context variation from assignment uncertainty. Neither context occupancy nor the present associations demonstrate biochemical interaction, metabolic activity or causal function.

Functional and intervention studies should now connect this ecological organization to what microbes do. Closely related strains can differ in metabolic capacity and host interaction^17,18^. Combining metagenomics, metatranscriptomics and metabolomics could test whether the same species differs in gene expression and metabolic activity across contexts. Longitudinal strain tracking could establish whether context changes occur while a strain persists within a person or accompany its replacement. Dietary and biotic studies can now test whether baseline context occupancy helps explain individual responses and whether beneficial responses accompany changes in the contexts of resident or introduced microbes. By making these settings measurable within each person, taxon-in-context provides a basis for understanding variable intervention responses and designing microbiome interventions around ecological individuality.

## METHODS

### Microbiome datasets

The primary dataset was obtained from the Curated Metagenomics Database^10^ and is referred to here as CMD. Samples were required to be stool samples from individuals annotated as healthy in the source metadata and to have available age information. We then excluded samples with any recorded antibiotic use, treatment history, smoking history, positive fecal occult blood test, history of mumps, alcohol intake, pregnancy or preterm birth. Following these exclusions, the dataset comprised 7,700 shotgun metagenomic gut microbiome samples spanning early to later life. Available metadata included age, sex, country, body mass index, delivery mode, feeding practice and other subject level variables where available. Species level relative abundance profiles were derived using MetaPhlAn3^19^ and used for the primary analysis. This was a metadata defined health associated reference collection; incomplete reporting of clinical history and exposures means that absence of an annotation did not establish absence of the corresponding condition or exposure.

To evaluate whether microbial contexts generalized beyond species level data, we constructed a larger genus-level compendium by combining the Gut Microbiome Reference^4^ and Human Microbiome Compendium^12^ cohorts and filtering for samples annotated as healthy. This dataset, referred to as GMR-Compendium, consisted of 9,860 healthy gut microbiome samples, including 5,230 samples from GMR and 4,630 samples from the Human Microbiome Compendium. This compendium partially overlaps with CMD; dataset construction is described in <u>Supplementary</u> <u>Methods 4</u>. Taxonomic profiles were harmonized across the two sources by matching genus names, retaining only genera that could be consistently identified in both datasets for downstream analysis. Abundance profiles were normalized within samples to relative abundance.

For further evaluation of CMD-derived contexts, we used an independent longitudinal rUTI cohort^11^. This dataset included stool metagenomic samples collected from diagnosis through post treatment recovery in 106 participants. The first UTI episode was used for the present analysis, spanning baseline through approximately 180 days post treatment. The cohort included 63 participants without recurrence and 43 participants with recurrence of UTI, for a total of 480 metagenomic samples. The number of contexts and their taxonomic distributions were held fixed; only sample-specific context activation and taxon-level context occupancy were inferred for the UTI samples. Neither UTI microbiome profiles nor recurrence outcomes contributed to learning or selecting the CMD context reference.

### Microbiome decomposition using Discovery

We developed Discovery, an ensemble-based implementation of Latent Dirichlet Allocation (LDA)^5^, to stabilize microbial context inference across independently initialized model fits, and used it to decompose microbiome profiles. We refer to these components as microbial contexts, defined as latent taxonomic configurations inferred from the whole-community composition.

Before model fitting, species-level relative abundance profiles from the CMD dataset were filtered according to taxon abundance and sample prevalence. Retained values were taken from the original relative abundance table without renormalization, multiplied by 1,000 and truncated to non-negative integer pseudo-counts. LDA was fitted in Python v3.9.16 using gensim.models.LdaModel in Gensim v4.3.3^20^. The mathematical formulation, threshold behavior, model priors and fitting parameters are provided in the <u>Supplementary Methods 1</u>.

### Discovery configuration and representative-model selection

Standard finite LDA requires the number of latent contexts to be specified before fitting, and independently initialized fits under identical settings can yield different context compositions^5^. Discovery addresses these two aspects by selecting the number of microbial contexts and preprocessing thresholds, and by identifying a complete LDA fit representative of the recurrent context structures recovered across independent initializations (Fig. 1a).

Each configuration specified the number of microbial contexts (*K*) and the taxon abundance and sample prevalence thresholds used during preprocessing. Candidate values of *K* ranged from two to ten, and both thresholds ranged from 0.001 to 0.005 in increments of 0.001. We fitted 500 LDA models per configuration using independent random initializations.

For each configuration, context estimates from the 500 fitted models were pooled and partitioned into *K* recurrent context clusters using spectral clustering^21,22^. Isolation forests^23^ identified outlier context estimates within each cluster. A complete LDA fit remained eligible for selection only if all its context estimates passed this screening. The representative model for each configuration was the eligible fit with the smallest mean Euclidean distance between its context estimates and the median centres of their corresponding clusters.

To compare configurations, we developed a cardinality corrected version of the silhouette score^24^, which assessed context cluster cohesion and separation while penalizing deviations from the expected cluster cardinality. We selected *K* by applying a stability breakpoint rule to the cardinality corrected silhouette score profile across candidate configurations (Supplementary Fig. 1). Within the selected *K*, we retained the configuration with the highest cardinality corrected silhouette score. Full details of clustering, outlier screening, representative model selection and configuration selection are provided in <u>Supplementary Methods 1.</u>

We separately characterized stochastic variation among individual LDA fits across the evaluated configurations (<u>Supplementary Methods 3</u>; Supplementary Fig. 6; Supplementary Table 8). Within the CMD configuration selected in the reference Discovery run, we further compared context compositions and taxon-in-context assignments across individual LDA fits and assessed their agreement with the representative model (<u>Supplementary Methods 3</u>; <u>Supplementary</u> <u>Results 2</u>; Supplementary Fig. 7).

To assess reproducibility of the complete Discovery procedure, we performed five additional CMD Discovery runs using independent, nonoverlapping ensembles of 500 LDA initialization seeds per configuration. The input dataset, candidate configuration grid, and preprocessing, fitting and selection procedures were held constant, while configuration selection was repeated in each run. We compared the selected configurations, context compositions and taxon-in-context assignments across the reference run and the five additional runs (<u>Supplementary Methods 3</u>; <u>Supplementary Results 2</u>; Supplementary Fig. 3).

### Microbial context activation and taxon-in-context assignment

The selected representative LDA model provided two complementary representations of each microbiome profile. First, each sample was represented by a microbial context activation profile. The activation of context was defined as its posterior mixture weight in a sample. The activation values were non-negative and summed to one across *K* contexts within a sample.

Second, each retained taxon represented by a positive pseudo-count in a sample was assigned to the microbial context with the greatest posterior probability for that sample-taxon observation (Fig. 1b). We refer to this sample-specific assignment as a taxon-in-context assignment. The posterior probability calculation and inference settings are provided in <u>Supplementary Methods</u> <u>1</u>.

Sample-taxon observations with a positive original relative abundance that were transformed to a pseudo-count of zero did not receive taxon-in-context assignment, i.e., no-context state. Taxa excluded during vocabulary filtering were outside the taxa retained for context inference. The no-context state was retained in analyses for which assignment availability was informative but was not treated as an additional microbial context when context entropy was calculated.

### Naming and characterization of microbial contexts

Each microbial context was characterized at two distinct levels. The taxon with the largest within context probability was used as the species anchor in the context name. This anchor identifies the most characteristic taxon in the learned distribution; it does not imply that the context is generated by that species alone, that every member co-occurs with it or that all members share one fixed function.

Second, an architectural descriptor was assigned post hoc to summarize the internal organization of each context across samples and life stages. Representative taxa were identified using routed mass share, defined as the proportion of the total abundance assigned to a given context within a life stage that was contributed by each taxon. Taxa were ranked independently within each context and across five condensed life stages, and the representative set for each context was defined as the union of the five highest-ranking taxa in each stage.

Pairwise organization within each representative set was characterized using within-context Jaccard similarity, raw-presence Jaccard similarity, Pearson correlation between life-stage routed-mass profiles, and peak-stage distance. The metrics were integrated into within-context evidence scores for Complementary, Alternative, and Successive relationships. Pairs without sufficiently strong or distinct evidence were classified as Mixed or unresolved.

All names were assigned after model fitting and did not influence context inference, taxon-in-context assignment, or downstream statistical analyses. Exact routed-mass calculations, life-stage definitions, pairwise scoring weights, classification thresholds, and implementation-specific handling rules are provided in <u>Supplementary Methods 2</u>. These descriptors summarize population-level ecological organization and do not establish direct cooperation, competition, within-person succession, or shared biochemical function.

### Quantifying ecological context flexibility with context entropy

Ecological context flexibility was quantified for taxa observed in more than one context across the dataset. Within each resampled age-study cell, the distribution of context assignments for taxon i was summarized using normalized Shannon entropy^25^, hereafter context entropy,

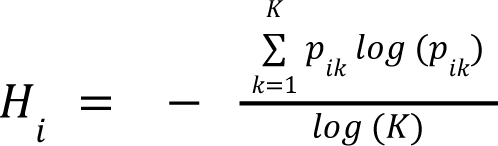

Where *p_ik_* is the proportion of assignments of taxon *i* to context *k*, and *K* is the number of learned microbial contexts (four in CMD and five in the GMR-Compendium). Normalization by *log*(*K*) constrained flexibility to a common 0-1 scale, allowing comparison across datasets. Taxon-level entropy values were averaged using taxon prevalence within the resampled subjects as weights, so that frequently observed taxa contributed more than sparsely detected taxa without directly weighting the estimate by their abundance.

To control for unequal study sizes, age-specific estimates were generated using study-balanced resampling. For each qualifying age-study cell, 15 participants were sampled in CMD and 12 in the GMR-Compendium, and one sample was selected per participant. The procedure was repeated 120 times and averaged within study, and the age-level estimate was the unweighted mean across contributing studies, so that large studies did not dominate the trajectory.

Because adolescent and young-adult ages had substantially weaker and less balanced support, the early-life and adult trajectories were fitted independently. Early life was defined as ages up to 3 years in both datasets; adulthood began at age 23 in CMD and age 18 in GMR-Compendium.

Primary-supported age estimates within each interval were smoothed using locally weighted linear regression^26^ with a LOWESS fraction of 0.30. Smoothing was performed on the signed square-root transformation of age to provide greater resolution during early life while reducing the leverage of widely spaced adult ages.

Uncertainty was quantified using 300 hierarchical bootstrap replicates^27^. Participants were resampled with replacement within each age-study cell, one sample was selected per resampled participant, and contributing studies were subsequently resampled with replacement within age. The complete age-specific curve was refitted for every bootstrap replicate, and pointwise 95% confidence bands were defined by the 2. 5^th^ and 97.5^tℎ^ percentiles of the bootstrap curves. Estimates from the lower-supported adolescent interval were calculated separately for diagnostic purposes but were not used to fit either supported trajectory.

The condensed life stage categories used in stage stratified analyses were Early Childhood, 0 to 4 years; Adolescent Transition, 4 to 16 years; Adulthood Transition, 16 to 33 years; Midlife Stability, 33 to 57 years; and Late life Transition, greater than 57 years. Lower age bounds were treated as inclusive and upper bounds as exclusive.

To complement context entropy with a compositional view, Fig. 2f displayed the seven taxa with the highest routed mass share in each context-life-stage combination. Within each context, the union of these taxa across life stages defined the common set displayed in the petal plots.

### Identification of age-associated context-responsive taxa and context pivots

To identify taxa whose assigned microbial context carried age related information beyond focal taxon abundance, we used a taxon specific nested regression framework. The reduced model included focal taxon abundance, represented using a cubic B-spline^28^ basis applied to log abundance. The full model included focal taxon abundance, together with microbial context assigned to the focal taxon, encoded as context identity indicator variables. For continuous age, ordinary least squares models^29^ were fitted and nested models were compared using partial F-tests. Model improvement was quantified as Δ*R*^2^ between the full and reduced models, and raw *P* values were corrected across eligible taxa using the Benjamini-Hochberg (BH) false-discovery-rate procedure^30^. The resulting BH-adjusted *P* values are denoted *q*. Taxa with *q* < 0. 01 and positive abundance-controlled Δ*R* were designated context-responsive taxa.

To identify assignment effects beyond both focal abundance and global context composition, a separate composition-controlled comparison was performed independently across all eligible taxa. The reduced model contained flexible focal-taxon abundance and the sample-level context activation profile, whereas the full model additionally contained the observed focal-taxon-in-context assignment. This effect was calibrated against a composition-preserving permutation null in which, for each sample, a null assignment was generated from its context activation probabilities restricted to the contexts retained for that taxon. Each null assignment was evaluated using the same composition-controlled reduced and full model contrast as the observed assignment. Empirical *P* values were calculated from 1,000 null realizations and BH-adjusted across all eligible taxa. The resulting BH-adjusted empirical *P* values are denoted*q_emp_*. Taxa with *q_emp_* < 0. 05 and positive observed composition-controlled 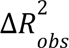 were designated context pivots. Full model specifications, filtering criteria, and permutation procedures are provided in <u>Supplementary Methods 5</u>.

### Delivery mode associated context redistribution

Delivery mode-associated context redistribution was analyzed in infants aged 0 to 1 year with available delivery mode metadata (n = 1650; 932 vaginally delivered [VD] and 718 delivered by cesarean section [CS]). For each focal taxon, we tested whether taxon-in-context assignment was associated with delivery mode after adjustment for species abundance. Multiple-testing correction was performed using the Benjamini–Hochberg procedure, with BH-adjusted *P*-values reported as *q*-values. For taxa significantly associated with delivery mode, context redistribution between delivery groups was quantified using Jensen Shannon divergence^31^ between context assignment distributions.

### Definition of typical and atypical microbiome configurations

Typical and atypical aging related microbiome configurations were defined from adulthood transition through late life. Early childhood samples were excluded from this analysis because the Successive E. coli context might be naturally prominent during early life^14,32^. Elevated activation of the Successive E. coli context was used to define atypical microbiome configuration. For each life stage, we calculated the mean activation of the Successive E. coli context across samples and used this value as the life-stage-specific activation threshold. Samples with activation above this threshold were classified as atypical. This classification yielded 4,278 typical samples and 744 atypical samples. We then tested whether taxon-in-context assignment was associated with typical versus atypical configuration after adjusting for taxon abundance. Activation contributes to both group definition and taxon assignment, therefore redistribution toward the Successive E. coli context is partly expected and does not independently establish an atypical aging state. Importantly, typical and atypical designations describe variations within a restricted health associated reference population and should not be interpreted as a contrast between healthy and diseased microbiomes.

### Ecological context switching

CMD-learned microbial contexts were projected onto the rUTI cohort after taxonomic harmonization. Participants were required to have at least two observed stool samples for within-participant switching analysis, yielding 105 eligible participants.

A taxon was considered to switch contexts within a participant when it received assignments to at least two different microbial contexts across that participant’s longitudinal samples. Across the full longitudinal series, 230 taxa received at least one valid context assignment and therefore constituted the taxon set evaluated for switching. This 230-taxon set differs from the 223-taxon feature set used in the subsequent recurrence-prediction analysis. For each participant, we quantified whether any taxon switched context, the number of switching taxa, and the number and identity of contexts occupied by each switching taxon. For transition-level analyses, strict context-switching events were defined as changes between two different microbial contexts for the same taxon between consecutive available samples from the same participant, with a valid context assignment required at both samples.

### UTI recurrence prediction

For recurrence prediction, we restricted the rUTI cohort described above to episode1 data, comprising 106 participants and 480 matched stool metagenomes. The prediction feature set was the 223 of 275 CMD taxa which was used consistently across all representations in the primary recurrence analysis.

Because recurrence could shorten longitudinal follow-up, later sample availability, visit count, missingness and follow-up duration could become associated with recurrence status and thereby introduce outcome-related sampling information. To reduce this source of leakage, samples were ordered by collection day within each participant and the primary analysis was restricted to the first two observed stool samples. Before ordering, missing collection days were imputed using the median for the corresponding scheduled timepoint; ties in collection day were resolved by scheduled timepoint and then sample identifier. Participants with fewer than two observed samples were excluded, yielding 105 participants and 210 metagenomes. The original scheduled study timepoint, total visit count, missingness, follow-up duration and availability of subsequent samples were not included as predictors; the retained samples were represented only by their within-participant ordinal position as the first or second observed sample. The distribution of these retained samples across the original study schedule and timing-only sensitivity analyses are described in <u>Supplementary Methods 6</u> and Supplementary Table 4. In the original cohort, recurrence ended the current episode and initiated a new episode^11^; restriction to episode 1 therefore supports interpreting these samples as preceding the recorded recurrence.

All representations in the primary recurrence analysis were constructed from the same 223 taxa at the same two ordinal samples for the same participants. The abundance representation used centred log ratio transformed^33^ abundance values for each taxon at each retained ordinal sample. The taxon-in-context assignment representation used one hot indicator encoding the CMD reference context assigned to each taxon at each retained ordinal sample. For taxon-sample pairs without a valid taxon-in-context assignment, all context indicators were set to zero. Because context assignment is conditional on a taxon meeting the 0.1% abundance threshold, we additionally evaluated a threshold-matched abundance-presence representation.

Recurrence status was encoded as the positive class. Prediction models used balanced L2 penalized logistic regression with the liblinear solver^34^. Model evaluation used fully nested cross validation: standardization and ANOVA F-test feature ranking were refitted within each inner training split, then refitted on the complete outer-training partition for outer-test prediction; the regularization strength and the number of retained features were selected using pooled inner out-of-fold (OOF) log loss, with AUROC used as a secondary criterion. The primary tuning grids were C = {0.03, 0.1, 0.3, 1, 3} and retained feature counts = {10, 20, 50, 100, 200, all}. Remaining tuning ties favoured fewer features and then smaller C. Evaluation used nested five-fold stratified cross-validation^35^, with three-fold inner cross-validation. Ninety-five percent CI were estimated using 2,000 patient-level bootstrap resamples of the outer-fold predictions. AUROC was calculated from pooled outer-fold predicted probabilities. Classification thresholds were selected exclusively from inner OOF training predictions by maximizing balanced accuracy and were locked before application to the corresponding outer test participants. Differences in AUROC between representations were calculated from paired participant-level OOF predictions, with CI obtained by the same participant-level bootstrap resampling. Consistency across classifier choices was additionally evaluated in a separate sensitivity analysis using five linear classifiers and a common 226-taxon feature universe across representations under repeated nested cross-validation (<u>Supplementary Methods 6</u>).

Feature stability was assessed across the outer cross-validation fits. A feature was considered stable if it was selected in at least 60% of outer fits and showed the same coefficient sign in at least 80% of its nonzero coefficients across selected fits. Median standardized coefficients were calculated across the fits in which the feature was selected.

## DATA AVAILABILITY

The primary analyses used publicly available microbiome datasets from the Curated Metagenomics Database, the Gut Microbiome Reference, the Human Microbiome Compendium and the longitudinal rUTI cohort of Choi et al. Original data and associated metadata are available from their respective repositories. The processed abundance matrices, sample manifests, taxonomic harmonization files, Discovery-derived context representations and data supporting the analyses reported in this study will be made available upon completion of peer review.

## CODE AVAILABILITY

The Discovery implementation and scripts used for preprocessing, model fitting, downstream statistical analyses and figure generation will be made available upon completion of peer review.

## ACKNOWLEDGEMENTS

We thank Elena Sandalova, Francisco Codoñer, Jolanda Lambert, Stephan Schuster, Sophie Swinkels, Qibo Sun, Monique van der Voet, Matthieu Pichaud for their valuable comments during the development of this work.

## AUTHOR CONTRIBUTIONS

J.G., N.A., F.C. and E.A. conceived and designed the study, analysed the data and contributed materials and analysis tools. J.G., N.A. and F.C. performed the computational experiments. C.L., E.A. and J.G. led the interpretation of the findings. F.S., S.B., T.Z., H.W., Y.C., C.G. and J.K. contributed to interpretation of the findings and substantive revision of the manuscript. J.G., N.A., C.L. and E.A. wrote the manuscript. All authors reviewed and edited the manuscript.

