## Supplementary material for "Taxon-in-context reveals ecological individuality in the human gut microbiome": Minutia.Discovery - Supplementary Information.pdf

Supplementary Methods

### Supplementary Methods 1. Discovery Algorithm

#### *Mathematical formulation of microbiome decomposition using LDA*

We developed Discovery to establish a reproducible reference for comparing taxon-in-context profiles across microbiomes. Independently initialized Latent Dirichlet Allocation (LDA)[5] fits can yield different microbial contexts and consequently different interpretations of the same taxon in the same sample. Discovery uses repeated fitting to identify recurrent context structures and select a representative model supported across the ensemble. For each candidate combination of context number and taxonomic filtering thresholds, it groups similar context estimates, screens atypical estimates and selects one complete fitted model whose contexts jointly lie closest to the recurrent cluster centres. Candidate configurations are ranked by context-cluster cohesion, separation and representation across the ensemble. The resulting reference model is then used to infer sample-level context mixtures and taxon-in-context profiles.

LDA is a probabilistic mixed-membership model originally developed for count-based document data[5]. When applied to microbiome data, samples correspond to documents, microbial taxa correspond to terms and transformed taxon abundances correspond to term counts. The model represents each sample as a mixture of latent microbial contexts and each microbial context as a probability distribution over taxa.

Throughout these Supplementary Methods, $N$ denotes the number of microbiome samples, $V$ the number of retained taxa, $K$ the candidate number of microbial contexts and $R$ the number of independently initialized LDA fits evaluated for a configuration. The indices $d$, $j$, $k$, $n$, $r$, and $i$ denote a sample, taxon, microbial context, pseudo-count unit, fitted model and spectral context cluster, respectively.

For sample $d$, the vector

$$\boldsymbol{\theta}_d = (\theta_{d1}, \ldots, \theta_{dK})$$

represents its context-mixture distribution, where

$$\theta_{dk} \geq 0 \ \forall k \text{ and } \sum_{k=1}^{K} \theta_{dk} = 1$$

For context $k$, the vector

$$\boldsymbol{\beta}_k = (\beta_{k1}, \ldots, \beta_{kV})$$

represents its taxonomic probability distribution, where

$$\beta_{kj} \geq 0 \text{ and } \sum_{j=1}^{V} \beta_{kj} = 1$$

Under the LDA generative formulation,

$$\boldsymbol{\theta}_d \sim Dirichlet(\boldsymbol{\alpha})$$

and

$$\boldsymbol{\beta}_k \sim Dirichlet(\boldsymbol{\eta})$$

where $\boldsymbol{\alpha}$ and $\boldsymbol{\eta}$ are the priors on the sample-context and context-taxon distributions, respectively. For each pseudo-count unit $n$ in sample $d$, an occurrence-level latent context assignment is drawn as

$$z_{dn} \sim Categorical(\boldsymbol{\theta}_d)$$

Conditional on $z_{dn}$, the observed taxon identity $w_{dn}$, indexed within the model vocabulary, is modelled as

$$w_{dn} \sim Categorical(\boldsymbol{\beta}_{z_{dn}})$$

The fitted model therefore provides a context-mixture distribution for each sample and a taxonomic probability distribution for each context. Posterior taxon-in-context assignments were subsequently derived from the fitted model as described below.

*Model assumptions and methodological rationale*

Discovery applies an ensemble-based configuration- and model-selection procedure to finite LDA[5]. The LDA model uses taxonomic abundance information. The mixed-membership formulation of LDA allows each microbiome to contain a mixture of microbial contexts and assigns each taxon to a context within that sample. Consequently, the same taxon can be associated with different contexts in different samples. Because $K$ must be specified before model fitting, Discovery evaluates a prespecified grid of candidate context numbers and preprocessing thresholds.

A consistent context reference is necessary to distinguish variation between microbiomes from variation introduced by model initialization. Discovery assumes that context structures repeatedly recovered across independent LDA initializations represent computationally stable solutions for the observed dataset. Stability was evaluated from the cohesion, separation and cardinality of clusters formed by the context-specific taxonomic distributions recovered across fitted models. These criteria were used to rank configurations within the prespecified search space and to select one representative complete LDA model.

*Overview of the Discovery workflow*

Discovery makes two distinct selection decisions: it selects a representative LDA model within each candidate configuration, then selects among configurations. A configuration specifies the number of microbial contexts and the taxon abundance and prevalence thresholds.

Selection within each configuration:

1. Fit LDA 500 times using independent random initializations.
2. Represent each context by its ten highest-probability taxa. Align these truncated vectors over the union of retained taxa across fits, with zeros elsewhere and without renormalization. Pool these vectors and group similar estimates into $K$ recurrent context clusters using spectral clustering[21,22].

3. Screen atypical estimates within each cluster using isolation forests[23]. A complete LDA fit remains eligible only if all its context estimates pass screening.
4. Select the eligible complete fit with the smallest mean distance between its context estimates and their corresponding cluster centres. Retain this fit as the representative model for that configuration, preserving its full taxon probability distributions.

Selection across configurations

5. Score each configuration using the cardinality-corrected silhouette criterion, which combines context-cluster cohesion, separation and cluster-size balance. Scores are calculated from all context estimates before outlier screening.
6. For each candidate $K$, summarize stability as the $90^{th}$ percentile of the cardinality-corrected silhouette scores across all evaluated abundance and prevalence threshold combinations.
7. Apply the stability-breakpoint rule to the resulting stability profile. Identify pairs of consecutive $K$ values within similar stability that are followed by a marked decline, with stability remaining below the pair's level for the next two higher-$K$ solutions. If multiple pairs qualify, retain the pair with the highest mean stability and select its larger $K$ as the stability-supported number of contexts
8. Among the configurations evaluated at the selected $K$, choose the configuration with the highest cardinality-corrected silhouette score and retain its representative model as the context reference for downstream inference.

This procedure preserves a complete fitted LDA model rather than constructing a reference by combining context estimates from different fits.

*Taxon filtering and pseudo-count transformation*

Before LDA fitting, species-level relative abundance profiles were filtered according to taxon abundance and prevalence. For each candidate Discovery configuration, taxon filtering was determined by an abundance threshold $\tau_a$ and a sample-prevalence threshold $\tau_p$. Let $x_{dj}$ denote the original relative abundance of taxon $j$ in sample $d$. A binary threshold-passing indicator was defined as

$$I_{dj}(\tau_a) = \left\{1,\ x_{dj} \geq \tau_a;\ 0,\ x_{dj} < \tau_a\right\}$$

The number of samples in which taxon $j$ met the abundance threshold was

$$n_j(\tau_a) = \sum_{d=1}^{N} I_{dj}(\tau_a)$$

Taxon $j$ was retained when

$$n_j(\tau_a) \geq floor\lfloor N\tau_p \rfloor$$

A cohort-wide filtering was performed to apply the same eligibility rule to every taxon, thus avoiding taxa that were supported across pooled populations but occurred unevenly among individual studies.

The retained taxa comprise the LDA model vocabulary. After taxon selection, retained taxa were extracted from the original relative abundance table. Retained relative abundances were multiplied by 1,000 and truncated to integer pseudocounts, giving a per sample assignment floor of 0.1% relative abundance among retained taxa. These pseudocounts do not represent sequencing read counts.

$$c_{dj} = floor\lfloor 1,000x_{dj} \rfloor$$

Thus, any relative abundance satisfying $0 < x_{dj} < 0.001$ was converted to $c_{dj} = 0$.

*LDA implementation and fitting parameters*

LDA was implemented in Python v3.9.16 using gensim.models.LdaModel in Gensim v4.3.3[20]. For each configuration, num_topics was set to $K$. In the initial ensemble, random_state was set to the fitted-model identifier $r \in \{0, ..., R - 1\}$, with $R = 500$.

The parameters explicitly supplied to LdaModel were: update_every=1, iterations=50, alpha='symmetric', chunksize=2, passes=3, and per_word_topics=True. Parameters not explicitly supplied retained the Gensim v4.3.3 defaults.

*Candidate configuration space and ensemble fitting*

A Discovery configuration was defined by three parameters: the number of microbial contexts $K$, abundance threshold $\tau_a$ and sample-prevalence threshold $\tau_p$. Candidate values were

$$K \in \{2, 3, 4, 5, 6, 7, 8, 9, 10\},$$

$$\tau_a \in \{0.001, 0.002, 0.003, 0.004, 0.005\},$$

and

$$\tau_p \in \{0.001, 0.002, 0.003, 0.004, 0.005\}.$$

Each configuration was fitted $R = 500$ times using independently initialized LDA fits. Each fitted model contributed $K$ context-specific taxonomic distributions, giving $RK = 500K$ context estimates per configuration.

*Spectral clustering of context estimates*

Let $\beta_{rk}$ denote the truncated taxonomic weight vector for context $k$ from fitted model r. Each vector retained the ten highest-probability taxa. Vectors were aligned over the union of these retained taxa across all contexts and fits within the configuration. For each configuration, the pooled set

$$\boldsymbol{B} = \{\boldsymbol{\beta}_{rk}: r = 0, ..., R - 1;\ k = 1, ..., K\}$$

contained $|\boldsymbol{B}| = RK$ context estimates. These estimates were partitioned into $K$ context-clusters using SpectralClustering in scikit-learn v1.5.2[21,22]. The implementation specified n_clusters=K, assign_labels='discretize' and random_state=0. The remaining parameters retained their scikit-learn v1.5.2 defaults: eigen_solver=None, n_components=None, n_init=10, gamma=1.0, affinity='rbf', n_neighbors=10, eigen_tol='auto', degree=3, coef0=1, kernel_params=None, n_jobs=None and verbose=False.

Let $c(r, k) \in \{1,..., K\}$ denote the spectral cluster assigned to context estimate $\beta_{rk}$, and let

$$C_i = \{\boldsymbol{\beta}_{rk} \in \boldsymbol{B}\colon c(r, k) = i\}$$

denote spectral context-cluster $i$. Each $C_i$ comprised recurrent estimates of a latent microbial context across independently initialized LDA fits.

*Isolation-forest screening of context estimates*

An isolation forest identified outlier context estimates within each context-cluster. Specifically, an isolation forest was fitted separately to the context probability vectors within each context-cluster using IsolationForest in scikit-learn v1.5.2[23]. The parameters were n_estimators=50, max_samples='auto', contamination=0.1, max_features=1.0 and random_state=42; remaining parameters retained their scikit-learn v1.5.2 defaults.

*Eligible LDA fits and context-cluster centers*

A fitted model $r$ was eligible for representative-model selection when all its $K$ context estimates remained after isolation-forest screening. The center of context-cluster $C_i$ was defined as the component-wise median of the taxonomic probability vectors assigned to that cluster. For taxon $j$
,

$$\tilde{\mu}_{ij} = median_{\beta_{rk \in C_i}}(\boldsymbol{\beta}_{rkj}).$$

Thus, the vector

$$\tilde{\mu}_i = (\tilde{\mu}_{i1}, ..., \tilde{\mu}_{iV})$$

contained the median probability of each taxon across all context estimates assigned to $C_i$.

*Selection of a complete representative LDA model*

For each eligible fitted model $r$, the Euclidean distance between context estimate $\boldsymbol{\beta}_{rk}$ and the center of its assigned spectral cluster was

$$\delta_{rk} = \left|\boldsymbol{\beta}_{rk} - \tilde{\mu}_{c(r,k)}\right|_2$$

The mean distance across the $K$ context estimates from fitted model $r$ was

$$\bar{\delta}_r = \frac{1}{K}\sum_{k=1}^{K} \delta_{rk}$$

The eligible fit with the smallest $\bar{\delta}_r$ was selected as the representative model for that configuration. This procedure selected one complete observed LDA fit rather than averaging or combining contexts obtained from different fits. It therefore preserved the correspondence among the context-specific taxonomic distributions, sample-specific context mixtures and per-taxon posterior context probabilities estimated within the same LDA realization.

*Cardinality-corrected silhouette score*

Configuration stability was evaluated using the cohesion, separation and cardinality of the spectral context clusters. Let $m_i = \left|C_i\right|$ denote the cardinality of cluster $C_i$. Because each of the $R = 500$ fitted models contributed $K$ context estimates, the expected cardinality of each cluster was

$$E = \frac{|\boldsymbol{B}|}{K} = R = 500$$

For context estimate $u \in \beta$, the conventional silhouette[24] value was calculated using Euclidean distance:

$$s(u) = \frac{b_{near}(u) - a_{in}(u)}{max\{a_{in}(u), b_{near}(u)\}},$$

where $a_{in}(u)$ is the mean distance between $u$ and the other estimates in its own cluster and $b_{near}(u)$ is the smallest mean distance between $u$ and the estimates in any other cluster. The mean silhouette value of cluster $C_i$ was

$$\bar{s}_i = \frac{1}{m_i} \sum_{u \in C_i} s(u)$$

Singleton observations received a silhouette value of zero under the scikit-learn implementation.

To avoid conflict with the LDA prior α, the cardinality factor is denoted by $\omega_i$:

$$\omega_i = \{\frac{m_i}{E}, m_i \leq E;\ 1 - \frac{m_i - E}{E} = 2 - \frac{m_i}{E}, m_i > E$$

The cardinality-corrected score for cluster $C_i$ was

$$S_i^{corr} = \omega_i \bar{s}_i.$$

The configuration-level corrected silhouette score was the unweighted arithmetic mean across the $K$ clusters:

$$S_{config} = \frac{1}{K} \sum_{i=1}^{K} S_i^{corr}$$

No lower bound was imposed on $\omega_i$ in the implementation, allowing it to take negative values for sufficiently oversized clusters. Future implementations should consider bounding this factor at zero to improve the interpretability and numerical behavior of the corrected silhouette score.

Silhouette and cardinality scores were calculated using all context estimates before isolation-forest screening.

*Selection of the number of microbial contexts based on stability-breakpoint rule*

To select the number of microbial contexts ($K$), we applied a stability-breakpoint rule to the cardinality-corrected silhouette scores from the complete configuration grid.

For each candidate number of contexts $K$, stability was summarized as

$$S_K = Q_{0.90}(s_{K,1},..., s_{K,n_K})$$

where $s_{K,i}$ is the cardinality-corrected silhouette score for candidate configuration $i$ at $K$, $n_K$ is the number of configurations evaluated at that $K$, and $Q_{0.90}$ denotes the $90^{th}$ percentile. This robust upper-tail summary reduced dependence on a single unusually high-scoring configuration while retaining information about the best-performing region of the search space.

Each pair of consecutive context numbers, $K - 1$ and $K$, was evaluated as a potential local stability plateau. The pair qualified as locally stable when

$$\left|S_K - S_{K-1}\right| \leq \delta$$

where the adjacency tolerance was prespecified as $\delta = 0.05$. The stability level of the pair was defined as

$$P_K = \frac{S_{K-1} + S_K}{2}$$

The larger value $K$ was considered a candidate stability breakpoint when the immediately following solution showed a decline of at least $\Delta = 0.10$:

$$P_K - S_{K+1} \geq \Delta$$

To prevent selection based on an isolated decrease, the decline was additionally required to persist at the next higher context number:

$$S_{K+2} < P_K$$

When multiple candidate breakpoints satisfied these criteria, the candidate associated with the largest plateau stability $P_K$ was selected. If plateau stability was tied, the larger $K$ was retained. Within the $K$ selected by the stability breakpoint rule, the final abundance and prevalence threshold combination was the configuration with the highest cardinality-corrected silhouette score. The resulting stability profiles and identified breakpoints are shown in Supplementary Fig. 1.

*Microbial context activation*

For the representative LDA model, variational inference estimated the Dirichlet parameter vector

$$\boldsymbol{\gamma}_d = (\gamma_{d1},\dots,\gamma_{dK})$$

for each sample. The posterior sample-level context activation was obtained by normalizing these parameters:

$$\hat{\theta}_{dk} = \frac{\gamma_{dk}}{\sum_{l=1}^{K}\gamma_{dl}}$$

Thus,

$$\hat{\theta}_{dk} \geq 0 \; and \; \sum_{l=1}^{K} \hat{\theta}_{dk} = 1.$$

The vector $\hat{\theta}_d$ was termed the microbial context activation profile of sample $d$. Activation values represent posterior mixture weights rather than binary indicators of context presence.

<u>*Taxon-in-context assignment*</u>

Taxon-level context probabilities were obtained using the Gensim get_document_topics function with per_word_topics=True and minimum_probability=0.0. For taxon $j$ represented by a positive pseudo-count in sample $d$, the variational probability of context $k$ was

$$\phi_{djk} = \frac{exp\{E_q[log\theta_{dk}] + E_q[log\beta_{kj}]\}}{\sum_{l=1}^{K} exp\{E_q[log\theta_{dl}] + E_q[log\beta_{lj}]\}},$$

where $q$ denotes the variational posterior distribution. Under this posterior,

$$E_q[log\theta_{dk}] = \psi(\gamma_{dk}) - \psi(\sum_{l=1}^{K} \gamma_{dl}) ,$$

$$E_q[log\beta_{kj}] = \psi(\lambda_{kj}) - \psi(\sum_{v=1}^{V} \lambda_{kv}).$$

where $\psi$ is the digamma function, $\gamma_d$ contains the variational parameters of the sample-context distribution and $\lambda_k$ contains the variational parameters of the context-taxon distribution.

Gensim reports per-taxon relevance values after multiplying $\phi_{djk}$ by the transformed abundance weight $c_{dj}$:

$$\rho_{djk} = c_{dj}\phi_{djk}.$$

For a fixed sample-taxon observation with $c_{dj} > 0$, the same value of $c_{dj}$ is applied across contexts. Therefore,

$$argmax\rho_{djk} = argmax\phi_{djk}.$$

To distinguish the derived taxon-level assignment from the occurrence-level latent variable $z_{dn}$ in the generative model, the taxon-in-context assignment is denoted by

$$\hat{z}_{dj} = argmax\phi_{djk}.$$

The model retains probabilistic information across contexts during inference; hard assignment provides an interpretable categorical representation for downstream analyses. This simplification does not propagate assignment uncertainty into those analyses, but provides a consistent operational definition of a taxon's dominant context in each sample. Setting minimum_probability=0.0 prevented contexts from being omitted from the returned sample-level context distribution. No additional abundance, posterior-probability or assignment-confidence criterion was applied when accepting the maximum-probability assignment beyond taxon filtering and pseudo-count transformation. Consequently, every retained taxon represented by a positive pseudo-count in a sample received a taxon-in-context assignment.

Sample-taxon observations with a positive original relative abundance that were transformed to a pseudo-count of zero did not receive taxon-in-context assignment, i.e., no-context state. The no-context state was retained as a distinct assignment state in analyses for which assignment availability was informative, including the age-specific transition networks and urinary tract infection cohort. It was excluded from the set of microbial contexts when calculating context entropy for ecological context flexibility.

### **Supplementary Methods 2. Quantitative characterization of microbial-context architecture**

#### *Context anchors and post hoc architectural descriptors*

The internal ecological architecture of each microbial context was characterized after model fitting and selecting the representative LDA model. This characterization was descriptive and did not alter the four-context solution, sample-level context activation values, taxon-in-context assignments, or any downstream statistical analysis.

For context $(k)$ and taxon $(t)$, the fitted LDA parameter $(P(t \mid k))$ represents the probability of taxon $(t)$ within the population-level taxonomic distribution of context. This parameter quantity is a model parameter and is distinct from the relative abundance of the taxon in an individual microbiome sample. The taxon with the largest value of $(P(t \mid k))$ was used as the species anchor in the context name. The anchor was intended as a mnemonic identifier for the most characteristic taxon in the learned distribution and does not imply that the context is generated by that taxon alone, that all context members co-occur with it, or that they share a fixed biological function.

The architectural adjective was assigned separately and post hoc using the concentration and persistence of representative contributors across life stages together with their pairwise ecological organization. Context-level adjectives were therefore interpretative summaries of the representative-taxon sets and relationship matrices. Thus, neither the species anchor nor the architectural descriptor contributed to context inference.

*Representative taxa and routed mass share*

The analysis combined sample-level taxon-in-context assignments, species-level relative-abundance profiles, and sample metadata containing age. Samples with missing age were excluded from the routed-mass and life-stage profile calculations. The two sample-level Jaccard metrics were calculated using all samples available in their respective assignment and abundance inputs and were not restricted to samples with age metadata.

Samples with available age were assigned to five condensed life stages: Early Childhood, $0 \leq age < 4\, years$; Adolescent Transition, $4 \leq age < 16$; Adulthood Transition, $16 \leq age < 33$; Midlife Stability, $33 \leq age < 57$; and Late-life Transition, $age \geq 57$. These age groups captured broad developmental periods while retaining enough samples for analyses of microbial contexts. Their boundaries provided a consistent basis for comparing age related patterns across taxa and contexts, without implying distinct biological transitions.

Representative taxa were identified using the same procedure for all four contexts. For taxon (jt), context $(k)$, and life stage $(g)$, routed mass share was defined as

$$R\,(t,\ k,\ g)\ =\ \frac{\sum_{s\in Sg} A(s,t)\,\mathbb{1}\{Z(s,t)=k\}}{\sum_{s\in Sg}\sum_{u\in \boldsymbol{\mathcal{V}}} A(s,u)\,\mathbb{1}\{Z(s,u)=k\}}$$

where $A(s,t)$ is the relative abundance of taxon $(t)$ in sample $(s)$, $(Z(s,t))$ is its sample-level context assignment, $\mathbb{1}\{\cdot\}$ is an indicator function, $S_g$ as the samples in life stage $g$, and $\boldsymbol{\mathcal{V}}$ as the taxon set. The numerator therefore represents the pooled abundance mass contributed by taxon $(t)$ to context $(k)$ within life stage $(g)$, while the denominator represents the total abundance mass assigned to that context in the same life stage. Routed mass share consequently measures the fraction of a context's pooled assigned abundance contributed by a given taxon after sample-level context routing.

Within each context and life stage, taxa were ranked by descending routed mass share. The five highest-ranked taxa were retained, or all contributing taxa if fewer than five were available. The representative set for each context was the union of these taxa across the five life stages. The representative taxon set for context $k$ was defined as

$$T_k\ =\ \cup_5^{g=1} Top5_j\{R(t,k,g)\}$$

Applying the same selection rule to all contexts yielded representative sets of nine taxa for the *Bacteroides vulgatus* context, five for the *Bifidobacterium longum* context, ten for the *Escherichia coli* context, and seven for the *Prevotella copri* context.

The *B. longum* representative set remained at five taxa, the minimum possible union size under this selection rule, because the same five taxa repeatedly occupied the five highest routed-mass positions across life stages. This recurrent, compact contributor set provided the primary quantitative basis for the descriptor Concentrated. The relatively steep head of the corresponding within-context taxon probability distribution provided additional qualitative support for this interpretation.

<u>*Pairwise ecological-architecture metrics*</u>

Pairwise organization was evaluated for every unordered taxon pair $(i, j)$ within $T_k$ using four metrics.

*Within-context Jaccard similarity*

Let $S(i, k)$ denote the set of unique samples in which taxon $i$ was assigned to context $k$. Shared occupancy of the focal context was calculated as

$$J_{context}(i, j, k) = \frac{|S(i, k) \cap S(j, k)|}{|S(i, k) \cup S(j, k)|}$$

A value of one indicates that the taxa were assigned to the focal context in the same samples, whereas a value of zero indicates that they had no shared assigned samples.

*Raw-presence Jaccard similarity*

Let $P(i)$ denote the set of unique samples in which taxon $i$ had relative abundance greater than zero, irrespective of context assignment. General sample-level co-occurrence was calculated as

$$J_{presence}(i,\ j)\ =\ \frac{|P(i) \cap P(j)|}{|P(i) \cup P(j)|}$$

*Life-stage profile correlation*

Each selected taxon was represented by its five-element routed-mass-share profile,

$$R_{t,k} = \left[R(t,\ k,\ g_1),\ldots, R(t,\ k,\ g_5)\right]$$

A value of zero was used when a taxon had no routed mass in a particular life stage. Similarity between the profiles of taxa $i$ and $j$ was calculated using Pearson correlation:

$$r(i, j, k)\ =\ cor(R_{i,k},\ R_{j,k})$$

If either profile was constant across the five life stages, or if the calculated correlation was undefined, the correlation was set to zero. This metric quantified whether two taxa changed their relative contributions to the same context in parallel across life stages.

*Peak-stage distance*

The peak stage of each taxon was defined as the ordinal position of the maximum value in its routed-mass-share profile. Peak-stage distance was calculated as

$$D_{peak}(i, j, k)\ =\ |arg\ max\ g\ R(i, k, g)\ -\ arg\ max\ g\ R(j, k, g)|$$

When a profile had equal maxima in multiple stages, the earliest stage in the predefined life-stage order was selected, consistent with the implementation of `argmax`. If either profile had zero routed mass across all five stages, the distance was set to five and was subsequently treated as maximally separated.

Together, the two Jaccard metrics distinguished shared occupancy of the same latent context from general taxon co-occurrence, whereas the life-stage metrics distinguished aligned from temporally displaced participation. Abundance magnitude contributed to the life-stage analysis only through routed mass after sample-level context assignment and within-context, within-stage normalization.

*Within-context ranking and transformed evidence components*

Because the distributions of the pairwise metrics differed among contexts, scoring was based on empirical percentile ranks calculated separately within each context. All selected-taxon pairs, including pairs subsequently marked as low support, contributed to these within-context rank distributions.

For a nonconstant metric $x$ evaluated over $n_k$ pairs in context $k$, the high-direction rank was calculated as

$$H(x_i) = \frac{rankaverage(x_i)}{n_k}$$

where tied observations received their average rank. The resulting ranks ranged from 1/n□ to 1 and therefore did not necessarily include zero. When a metric had the same value for every pair in a context, all pairs were assigned a neutral rank of 0.5. For the two Jaccard metrics, evidence in the low direction was defined as

$$L(x) = 1 - H(x)$$

The positive and negative components of the life-stage profile correlation were defined as

$$r^{+} = max(r, 0) \text{ and } r^{-} = max(-r, 0)$$

and were percentile-ranked separately within each context.

Differences in the life stages at which two taxa peaked were summarized using complementary closeness and separation scores:

$$C_{peak} = 1.00\ if\ D_{peak} = 0;\ 0.75\ if\ D_{peak} = 1; 0.00\ if\ D_{peak} \geq 2\ .$$

$$S_{peak} = 0.00\ if\ D_{peak} = 0;\ 0.35\ if\ D_{peak} = 1; 1.00\ if\ D_{peak} \geq 2\ .$$

Because peak-stage distance was derived from only five broad life-stage categories, it was treated as an ordinal. Peaks occurring in the same stage were considered strongly aligned, whereas peaks in adjacent stages were treated as approximately aligned to tolerate discretization and boundary effects. A separation of at least two stages was considered stronger evidence of temporal displacement. The assigned values were operational coding choices used to preserve these ordered categories and were not estimated from the data. Because both components were subsequently percentile-ranked independently within each context, their absolute numerical magnitudes did not directly determine the final evidence scores.

*Pairwise evidence scores*

Three evidence scores were calculated for every selected-taxon pair:

$$E_{comp} = 0.35H(J_{context}) + 0.15H(J_{presence}) + 0.35H(r_{+}) + 0.15H(C_{peak})$$

$$E_{alt} = 0.35L(J_{context}) + 0.30L(J_{presence}) + 0.20H(r_{-}) + 0.15H(C_{peak})$$

$$E_{succ} = 0.25L(J_{context}) + 0.10L(J_{presence}) + 0.25H(r_{-}) + 0.40H(S_{peak})$$

The weights were fixed operationally to reflect the defining evidence for each relationship class and summed to one within each score. Complementary evidence assigned the greatest weight to shared context occupancy and positively aligned life-stage profiles, with general co-presence and peak-stage proximity providing supporting evidence. Alternative evidence was driven primarily by weak context co-occupancy and weak general co-presence, while nearby peak stages and negative profile correlation provided secondary evidence distinguishing alternative organization from temporal succession. Successive evidence assigned the greatest weight to peak-stage separation, supported by weak simultaneous context occupancy and negative life-stage profile correlation; general co-presence received a smaller weight because taxa participating at different life stages could nevertheless occur in some of the same samples.

The transformations and weights were rule-based analytical choices and were not optimized against a clinical or ecological outcome. The resulting scores quantify relative evidence among selected taxon pairs within each context and should not be interpreted as probabilities of ecological interaction.

*Support assessment and pairwise classification*

Support was assessed using the numbers of unique samples in the assignment union and raw-presence union. A pair was marked as Low support when

$$|S(i,k)\ \cup\ S(j,k)| < 3 \text{ or } |P(i)\ \cup\ P(j)| < 3$$

The reported analysis used `min-support = 3`. Evidence scores were calculated for all pairs to retain a common within-context ranking distribution, but pairs meeting either low-support condition were reported as Low support and were not assigned an ecological relationship class.

For each supported pair, let $E(1)$ and $E(2)$ denote the largest and second-largest of $E_{comp}$, $E_{alt}$, and $E_{succ}$. Their separation was defined as

$$\Delta E\ =\ E(1)\ -\ E(2).$$

The class corresponding to $E(1)$ was treated as the dominant relationship. Pairwise labels were assigned as follows:

- Mixed or unresolved when $E(1)\ <\ 0.52$ or $\Delta E\ <\ 0.08$;
- a leaning Complementary, Alternative, or Successive relationship when $E(1)\ \geq\ 0.52$ and $0.08\ \leq\ \Delta E\ <\ 0.16$; and
- a strong Complementary, Alternative, or Successive relationship when $E(1)\geq 0.52$ an $\Delta E \geq 0.16$.

For visualization, strong relationships were represented by uppercase codes (`C`, `A`, or `S`), leaning relationships by lowercase codes (`c`, `a`, or `s`), Mixed relationships by `M`, and Low-support pairs by

?. Strong and leaning calls were combined within their corresponding parent classes when descriptively summarizing a relationship matrix.

*Context-level interpretation*

The pairwise procedure classified relationships between representative taxa; it did not automatically assign a single architectural class to an entire context. The final architectural adjectives were assigned post hoc as concise descriptive summaries of the representative-taxon structure and the corresponding pairwise relationship matrix. No additional percentage threshold, hypothesis test, or trained classifier was used to convert the pairwise calls into context-level names.

Concentrated referred primarily to the persistence of a compact representative contributor set across life stages. Mixed, Complementary, and Successive summarized the characteristic organization observed in their respective pairwise matrices. The reuse of pairwise relationship terms in the context names should therefore not be interpreted as an additional formal context-level classification.

Complementary pairs tended to occupy the same context in the same samples and to show aligned life-stage contribution profiles. Alternative pairs tended to show weak context co-occupancy and weak general co-presence without sufficient peak-stage separation to support a successive pattern. Successive pairs tended to show weak simultaneous context occupancy together with negatively aligned or temporally displaced contribution profiles. Mixed or unresolved pairs lacked sufficiently strong or distinct evidence for any of these three relationships.

These labels are operational descriptions of population-level ecological architecture. They do not demonstrate direct mutualism, competition, functional equivalence, biochemical interaction, or sequential replacement of taxa within individual participants.

### Supplementary Methods 3. Analysis of stochastic variability and reproducibility of Discovery

#### *Across-configuration-space analysis of stochastic LDA variability*

This analysis characterized variation among independently initialized LDA fits before applying the Discovery stabilization procedure. We evaluated configurations that varied in the number of microbial contexts and the abundance and sample prevalence thresholds. A corresponding no-filter configuration, with both thresholds equal to zero, was also included.

Repeated LDA models were fitted independently under each configuration using different random initialization seeds. A complete realization comprised the full set of $K$ context-specific taxon-probability distributions produced by one LDA fit. The sampled realizations were used for both the $K$-specific visualization of context-composition landscapes (Supplementary Fig. 6) and the quantitative fit-to-fit comparison (Supplementary Table 8).

*Within-selected-Discovery configuration analysis of stochastic LDA-fit stability*

To characterize stochastic variation among individual LDA realizations and evaluate the stabilization procedure used by Discovery, we performed a Within-selected-Discovery configuration analysis using the LDA fits generated under the configuration selected in the reference Discovery experiment. The selected Discovery configuration consisted of four microbial contexts, 0.003 abundance threshold, and 0.005 sample-prevalence threshold.

*LDA realizations and Isolation Forest classification*

All LDA realizations generated under the selected Discovery configuration were considered. Because the dataset, preprocessing, number of microbial contexts and LDA hyperparameters were held constant, variation among these realizations arose from differences in random initialization.

Discovery identifies recurrent context estimates across these stochastic realizations and applies Isolation Forest screening to identify atypical context solutions before selecting the representative realization. For the present analysis, LDA fits with a complete set of context estimates that remained after Isolation Forest screening were classified as eligible models, whereas fits identified during screening were classified as outliers. The Discovery-selected representative LDA realization was considered separately from both groups in visual summaries.

The purpose of this analysis is to characterize the extent of stochastic variation among individual LDA realizations and to determine whether the stabilization procedure preferentially retained the recurrent solution family represented by the selected model.

*Alignment of context compositions*

Each microbial context was represented by its complete taxon-probability distribution from the corresponding LDA realization. Context labels are arbitrary across independently initialized LDA fits; therefore, contexts from each realization were aligned to the representative architecture before comparison.

For each complete LDA realization, Jensen-Shannon distances[31] were calculated between all estimated contexts and all contexts of the representative model using the full taxonomic probability distributions. A one-to-one correspondence was then determined using the Hungarian algorithm[36] to minimize the Jensen-Shannon distance between the realization and the representative architecture. Context identities used for subsequent analyses were inherited from the corresponding representative contexts after alignment and therefore did not influence the alignment itself.

*Visualization of compositional variation among LDA realizations*

To visualize the overall compositional landscape of the individual LDA realizations, all complete context estimates were combined and pairwise Jensen-Shannon distances were calculated across their full taxonomic probability distributions. Principal coordinate analysis was then performed on this distance matrix[37]. This visualization was used to assess whether individual LDA realizations predominantly converged into recurrent context-specific compositional regions and whether outlier realizations showed greater displacement from these recurrent structures.

*Compositional distance from the representative architecture*

Compositional stability was quantified using the Jensen-Shannon distance between each aligned context estimate and its corresponding context in the representative realization. A Jensen-Shannon distance of zero indicates identical probability distributions, whereas increasing values indicate progressively greater compositional dissimilarity.

Distances were summarized separately for eligible models and outliers and stratified by representative-aligned context identity. This comparison quantified the degree to which the context estimates retained by Discovery remained close to the representative architecture and whether Isolation Forest outliers contained a broader range of compositional deviations.

*Sample-taxon assignment stability*

To determine whether differences among stochastic LDA realizations propagated to the individualized output of Discovery, each complete LDA realization was also compared with the representative model at the sample-taxon assignment level. Context assignments were first mapped to the representative context identities using the architecture-derived correspondence described above.

For each non-representative LDA realization, exact assignment agreement was calculated over sample-taxon observations present in both that realization and the representative model:

$$Exact\ agreement\ =\ \frac{number\ of\ sample{-}taxon\ observation\ assigned\ to\ the\ same\ aligned\ context}{number\ of\ sample{-}taxon\ observations\ compared}$$

Agreement was expressed as a percentage and summarized separately for eligible models and outliers. This analysis assessed whether compositional departures from the recurrent LDA architecture resulted in material differences in downstream individualized taxon-in-context assignments.

*Between-Discovery trials reproducibility analysis*

*Independent Discovery experiments*

Reproducibility was assessed by repeating the complete Discovery workflow using independent sets of LDA random initializations. The dataset, taxonomic preprocessing, filtering procedure, candidate configuration space, LDA hyperparameters other than the random initialization, and model-selection criteria were held constant across experiments. The original Discovery experiment was designated the reference run, and five additional Discovery experiments were performed using independent, non-overlapping ensembles of 500 LDA random seeds. Thus, differences among experiments reflected stochastic variation in LDA fitting rather than changes in the input data or analytical procedure.

For each experiment, candidate Discovery configurations were evaluated across the predefined combinations of number of microbial contexts, minimum relative-abundance threshold and sample-prevalence threshold. Each candidate configuration was evaluated from multiple LDA realizations, recurrent context structures were identified, and a representative realization was selected using the same Discovery model-selection procedure applied in the primary analysis.

*Configuration-level reproducibility*

Configuration-level reproducibility was evaluated by comparing the configurations selected independently by the reference run and the five repeated Discovery experiments. The number of microbial contexts, abundance-filtering threshold, sample-prevalence threshold, number of retained taxa and configuration-selection score were recorded for each experiment. Concordance of these quantities was used to determine whether independent stochastic repetitions of the complete Discovery workflow converged on the same or closely related model configuration.

This analysis was performed before comparing individual microbial context identities and therefore assessed the stability of the overall Discovery model-selection procedure.

*Composition-level reproducibility*

To determine whether repeated Discovery experiments recovered equivalent microbial contexts, each context was represented by its full taxonomic probability distribution from the representative LDA realization. Because LDA context labels are arbitrary across independent fits, contexts from each trial were aligned to those of the reference run before comparison.

For every trial, pairwise Jensen-Shannon distances were calculated between all trial and reference context distributions after harmonizing the taxonomic feature space. Missing taxa were assigned probability zero and each context distribution was renormalized to sum to one. A one-to-one correspondence between trial and reference contexts was determined using the Hungarian algorithm[36] to minimize the total Jensen-Shannon distance across matched contexts.

Context names were inherited from the corresponding reference context after alignment and were not used during matching.

Composition-level reproducibility was quantified using all unique pairwise Jensen-Shannon distances among the context distributions. Comparisons were classified as within-context when both distributions represented the same reference-aligned context identity and between-context when they represented different identities.

The within-context distribution therefore measures how much a given context varies when Discovery is repeated, whereas the between-context distribution provides the compositional separation expected between genuinely different microbial context identities. This comparison provides a reference scale for interpreting stochastic within-context differences: a context was considered structurally reproducible when repeated realizations remained closer to one another than to alternative context identities.

Between-context distances were additionally stratified by the six possible pairs of reference-aligned context identities to determine whether any particular pair of distinct contexts showed substantial compositional overlap.

*Assignment-level reproducibility*

Assignment-level reproducibility was assessed after fixing context correspondence using the context matching based on taxonomic composition. Assignment data were therefore not used to determine the mapping of context labels between runs, avoiding optimization of the alignment according to the assignment outcome itself.

Each individualized assignment was indexed by the combination of sample identifier and taxon identifier. The primary analysis was restricted to sample-taxon observations with a valid context assignment in all six Discovery runs, providing a common denominator for all reproducibility comparisons.

For each sample-taxon observation, the number of runs assigning it to its most frequently selected context was calculated. Assignment consensus was summarized as: 6/6 agreement; 5/6 agreement; 4/6 agreement; and ≤3/6 agreement.

The proportion of observations with unanimous 6/6 agreement and strong ≥5/6 agreement was calculated overall and separately according to the consensus microbial context.

In addition, exact assignment agreement was calculated for each unique pair of Discovery runs. For each pair, exact agreement was defined as the proportion of common sample-taxon observations receiving the same reference-aligned context assignment in both runs. Reporting all pairwise comparisons allowed assessment of whether reproducibility was general across independent Discovery experiments rather than being driven by similarity to the designated reference run.

As a sensitivity analysis, the union of sample-taxon observations across all runs was also evaluated by representing observations absent from an individual run as No assignment. This analysis was used to determine whether high agreement in the primary intersection-based analysis could be explained by preferential exclusion of observations without assignments in one or more runs.

### **Supplementary Methods 4. Reproducibility across datasets and enterosignature benchmarking**

*Construction of the GMR + Human Microbiome Compendium validation dataset*

To assess whether microbial contexts and their age associated patterns could be recovered across taxonomic resolutions and sequencing approaches, we constructed a larger genus level compendium by combining the Gut Microbiome Reference (GMR)[4] and Human Microbiome Compendium[12]. The resulting dataset, referred to as GMR-Compendium, comprised 9,860 healthy gut microbiome samples: 5,230 from GMR and 4,630 from the Human Microbiome Compendium.

Whereas CMD comprised species level shotgun metagenomic profiles, GMR-Compendium combined shotgun metagenomic profiles from GMR with 16S rRNA profiles from the Human Microbiome Compendium, harmonized at genus level. Discovery learned contexts separately in each datasets, allowing us to assess whether comparable ecological organization and age associated patterns emerged across these different datasets and measurement approaches. Metadata reconciliation identified at least 275 samples shared with CMD, corresponding to 2.79% of the initial GMR-Compendium dataset. These comprised 239 sequencing accession matches from BackhedF_2015/BKHD and 36 sample identifier matches from RaymondF_2016/RAY2016, corroborated by age, sex, country and sampling day. Among these identified shared samples, 239 were from early childhood (0–4 years) and 36 were from adults aged 21–35 years.

The combined dataset was also constructed to improve age coverage across the lifespan, particularly in late-adolescent and young-adult age bins that were underrepresented in the individual cohorts. Low sample sizes in these age intervals can increase trajectory variance and bias inference of age-associated microbial context dynamics. Combining compatible cohorts provided more uniform sampling across age, enabling more reliable estimation of lifespan-associated context activation and ecological flexibility patterns.

Genus-level features were harmonized across datasets using exact taxonomic name matching. Only genera that could be consistently identified across the combined dataset were retained for downstream analysis. Abundance profiles were normalized within each sample to relative abundance so that genus-level values summed to one per sample.

*Discovery analysis of the combined genus-level dataset*

Discovery was independently applied to the GMR-Compendium dataset to test whether microbial contexts comparable to those observed in CMD could be recovered without using CMD-derived assignments. This analysis was designed to distinguish dataset-specific structures from reproducible microbial ecological configurations.

The GMR-Compendium analysis preceded standardization of the software’s filtering grid. To account for its substantially larger sample size relative to CMD, we explored a different set of abundance and prevalence thresholds across several search batches. The CMD-based grid was later adopted as the released software's default, while the reported GMR-Compendium results remain based on the original dataset-specific search.

Applying the stability-breakpoint rule to these original search results supports five microbial contexts $K = 5$. The selected reference model used for downstream analyses had an abundance threshold of 0.005 and a prevalence threshold of 0.008, and retained 74 genera.

The resulting genus-level microbial contexts were compared with the CMD-derived species-level contexts using three criteria: dominant taxa or corresponding higher-level taxonomic groups, taxonomic membership structure, and age-associated context activation trajectories. Because CMD and GMR-Compendium differed in both sequencing platform and taxonomic resolution, exact one-to-one taxonomic identity was not required for correspondence. Instead, corresponding contexts were interpreted based on ecological similarity, including shared dominant or related taxa and comparable lifespan dynamics.

*Conceptual benchmarking against enterosignatures*

As an additional conceptual benchmark, Discovery-derived microbial contexts were compared with previously described enterosignatures[4]. This comparison was used to assess whether Discovery recovered recognizable population-level microbiome structures previously identified using an independent matrix decomposition framework.

Taxonomic contribution profiles from microbial contexts and enterosignatures were harmonized to a common taxonomic feature space. Jensen-Shannon distance was then calculated between each microbial context and each enterosignature, with lower values indicating greater similarity in taxonomic composition. For each microbial context, the closest matching enterosignature was identified as the pair with the lowest Jensen-Shannon distance. Correspondence was further evaluated by comparing dominant taxa and high-contribution member taxa within each paired structure.

This analysis compared the composition of microbial contexts with enterosignatures, but the two frameworks encode microbial organization differently. Enterosignatures summarize microbiome composition as additive combinations of shared components, whereas Discovery represents each sample as a mixture of microbial contexts and additionally assigns taxa to contexts within individual samples. The benchmark's scope was limited to a comparison of population-level ecological structure recovered.

### Supplementary Methods 5. Identification of age-associated context-responsive taxa and context pivots

A nested regression model-comparison framework was used to identify context-responsive taxa and context pivots. Individual focal taxon was analyzed independently. A focal taxon was considered eligible for age-association analysis only when at least two context groups remained after excluding contexts supported by fewer than 20 samples. This filtering criterion was applied to ensure that context-specific estimates were based on sufficient observations and that the regression model could compare age associations across multiple context assignments. Additionally, the analysis was restricted to samples with complete age information, species-level relative abundance values, taxon-in-context assignments, and microbial context activation scores. This yielded 93 eligible taxa.

Focal abundance was represented using a cubic B-spline basis applied to log abundance. The spline basis contained four columns for each eligible taxon. This representation captures thresholds, plateaus, and curved abundance-age relationships, providing a substantially stronger and fairer abundance baseline.

To identify the broad context-responsive taxa, an abundance-controlled nested model comparison was performed. The reduced model contained only the focal abundance block, while the full model additionally contained observed context assignment indicators:

$$Reduced: Age \sim f(A)$$

$$Full: Age \sim f(A) + S$$

where $f(A)$ represented the flexible focal taxon abundance, and $S$ represented the context assignment indicators. The abundance-controlled contribution of context assignment was quantified as the increase in the total model $R^2$, referred to as abundance-controlled $\Delta R^2$. Nested models were compared with partial F-tests. Raw $P$ values from each nested comparison were corrected across taxa with the Benjamini-Hochberg (BH) FDR procedure. The resulting BH-adjusted $P$ values are denoted $q$. Context-responsive taxa was identified based on this criteria: $q < 0.01$ and positive abundance-controlled $\Delta R^2$.

Subsequently, to identify context pivots, an observed composition-controlled effect was defined by comparing a model containing flexible abundance and context composition with the same model plus observed focal-taxon assignment:

$$Reduced: Age \sim f(A) + C$$

$$Full: Age \sim f(A) + C + S$$

where $C$ represents the activation values across all contexts within a sample. This was performed independently for all eligible taxa. The effect statistic was the total-scale increase in $R^2_{obs}$: observed $\Delta R_{obs}^{\ 2} = R^2 full - R^2 reduced.$ Additionally, adjusted between-context age gap measured as the maximum model-adjusted difference in mean host age among retained context groups was reported. To verify that taxon-in-context assignment does not merely recapitulate the sample's global context composition, permutation null was performed. For each 1,000 null realizations, a composition-compatible assignment $S^*$ was generated from the sample's context activation values, restricted to the contexts retained for the focal taxon and renormalized across those contexts. Each null assignment was evaluated using the same composition-controlled model contrast as the observed assignment. Importantly, the observed and null effects were evaluated against the same abundance- and composition-controlled reduced model, ensuring that they represented directly comparable quantities. The empirical $P$ value ($P_{emp}$) used the plus-one correction[38] and was BH-adjusted across all 93 eligible taxa. Context pivots were identified based on this criteria: empirical BH-adjusted empirical P value $q_{emp}$ < 0.05 and positive observed composition-controlled $\Delta R^2_{obs}$.

### Supplementary Methods 6. Sensitivity analyses of recurrent UTI prediction

#### *Temporal admissibility of the two-visit design*

To assess whether the two-visit recurrence analysis could exploit differences in sample timing, we performed a temporal-admissibility analysis using only sampling-time information. For each participant, stool samples were ordered by actual collection day and the first two observed samples were retained. Participants with fewer than two observations were excluded. The original scheduled study timepoint and actual collection day were used only for this temporal sensitivity analysis and were not included as predictors in the microbiome models.

We first characterized the correspondence between the two retained ordinal samples and the original study timepoints. This analysis was used to determine whether the first and second observed samples were concentrated at early scheduled timepoints or whether later samples frequently entered the two-visit representation.

We then evaluated whether sampling time alone could discriminate recurrence status. Three timing representations were considered: actual collection days for the two retained samples and their interval, numeric study timepoints and their interval, and a combined representation containing both sets of variables. For each representation, recurrence was modeled using linear logistic regression, spline-based logistic regression, and ExtraTrees[21,39] to detect both linear and nonlinear timing effects. Predictions were evaluated out of fold at the participant level.

Because an AUROC below 0.5 can represent outcome separation in the reverse direction, timing discrimination was summarized using direction-free AUROC, defined as max(AUROC, 1 − AUROC). A pre-specified practical-equivalence margin of 0.57 was used to identify timing-related discrimination that was sufficiently close to chance to be considered negligible. This margin was chosen to remain below weak-discrimination range around AUROC ≈ 0.60, thereby avoiding dismissal of small but potentially meaningful timing signals. Timing was considered practically equivalent to chance only if the one-sided 95% upper confidence bound of the direction-free AUROC did not exceed 0.57. This analysis was intended to test for residual calendar-related outcome information after fixing the number of microbiome observations to two per participant; it does not establish temporal precedence relative to recurrence because exact recurrence dates were unavailable.

*Robustness of recurrence discrimination across linear classifiers*

To assess whether the relative performance of the microbiome representations depended on classifier choice, we performed a separate sensitivity analysis using the same 105 participants and first two observed episode 1 samples described in the main Methods. This analysis used a 226-taxon universe, comprising CMD reference taxa with non-zero abundance in at least one of the 210 retained samples. The same taxa were used across all representations and classifiers. This feature universe differs from the 223 taxa used in the primary recurrence analysis; performance estimates from the two analyses were therefore reported separately. CMD-derived contexts and taxon-in-context assignments remained fixed and were not relearned from the rUTI cohort.

For the supplementary CLR representation, abundances were normalized across the full input community, restricted to the 226 taxa and renormalized to sum to one within this subcomposition. A pseudocount of $10^{-6}$ was added before taking natural logarithms and subtracting the sample-specific mean across the 226 taxa. Thresholded presence was defined at relative abundance ≥0.001 (0.1%), using full-community relative abundances, matching the eligibility threshold for taxon-in-context assignment. Context features comprised four binary indicators per taxon and ordinal sample; all four indicators were zero when no assignment was available. The two ordinal samples were concatenated within participants. Sampling-time variables and hospital identifiers were not used as predictors.

We compared five linear classifiers: L2- and L1-penalized logistic regression, elastic-net logistic regression, a linear support vector machine and shrinkage linear discriminant analysis. Logistic regression and the linear support vector machine used balanced class weights. L2 and L1 logistic regression used the liblinear solver[34], whereas elastic-net logistic regression used saga[40]. Performance was evaluated using 20 repetitions of nested stratified five-fold cross-validation, with identical participant partitions across representations and classifiers within each repetition. As in the primary analysis, feature standardization and F-statistic ranking were performed within each inner training partition and refitted on the complete outer-training partition before outer-test prediction, with 10, 15, 20, 30 or 50 retained features considered during three-fold inner validation. Features were ranked by descending computed F score, with exact score ties resolved by ascending column index and undefined scores ranked last; near ties were not rounded or merged. Hyperparameters and feature count were selected by inner-validation log loss, with mean inner-fold AUROC used to break log-loss ties. Regularization strengths were $C = 0.01, 0.1, 0.3, 1$ for L2 and L1 logistic regression, $C = 0.1, 0.3, 1$ for elastic-net logistic regression (L1 ratio 0.5), and $C = 0.01, 0.1, 1$ for the linear support vector machine. Support vector machine decision margins were transformed using the logistic sigmoid for log-loss tuning and score averaging, without fitted probability calibration. Shrinkage linear discriminant analysis used automatic covariance shrinkage[41] with the lsqr solver and tuned only the retained feature count.

Each participant received one held-out prediction per repetition. Primary AUROCs were calculated after averaging the 20 held-out predictions per participant. The arithmetic mean and range of repetition-specific AUROCs were calculated separately; these are distinct from AUROC calculated from averaged scores. Paired differences between taxon-in-context and each comparator were evaluated using 2,000 unstratified participant-level percentile bootstrap resamples of these averaged scores, with identical resampling indices across representations and classifiers. The resulting CI condition on the fitted predictions and were not adjusted for multiple comparisons.

As a complementary sensitivity analysis, recurrence labels were permuted 499 times using unrestricted participant-level shuffles, with the same permutation applied across representations and classifiers. The complete nested fitting procedure was repeated using five outer-validation repetitions for each permutation, including training-only standardization, feature ranking, hyperparameter selection and refitting. Stratified partitions were regenerated for each permuted outcome vector and shared across representations and classifiers. The unpermuted reference statistic was calculated using the same five-repetition procedure. The test statistic was the AUROC difference between taxon-in-context and the corresponding comparator, calculated from participant-level scores averaged across those five repetitions. One-sided permutation $P$ values were calculated using the plus-one correction[38]. Holm correction[42] [Holm, 1979] was then applied separately to the five CLR-abundance and five thresholded-presence comparisons, with the resulting Holm-adjusted values reported as $P_{adj}$.

Supplementary Results

### Supplementary Results 1. Internal ecological architecture of microbial contexts

The same representative-taxon selection procedure was applied to all four microbial contexts. Taking the union of the five taxa with the highest routed mass share within each of the five condensed life stages produced representative sets of nine taxa for the *B. vulgatus* context, five for *B. longum*, ten for *E. coli*, and seven for *P. copri*. These set sizes and the resulting pairwise classifications provided the quantitative basis for the post hoc architectural descriptors assigned to the four contexts. Representative-taxon contributions, pairwise metrics and relationship classifications are summarized in Supplementary Fig. 2.

#### *Complementary Prevotella copri context*

The *P. copri* context showed the strongest shared organization among the four microbial contexts. Its seven representative taxa generated 21 pairwise comparisons, of which 11 were classified as complementary or complementary-leaning, four as alternative, and six as successive; none were mixed or unresolved. Thus, more than half of the evaluated relationships supported shared context occupancy together with aligned life-stage contribution profiles.

This pattern provided the basis for the Complementary descriptor. The term denotes the comparatively strong tendency of representative taxa to participate in the context in the same samples and to exhibit parallel changes in routed mass share across life stages. It should not be interpreted as evidence of direct metabolic cooperation or mutualism.

*Successive Escherichia coli context*

The *E. coli* context showed the strongest temporally offset architecture. Its ten representative taxa generated 45 pairwise comparisons, comprising 22 successive or successive-leaning relationships, 19 alternative relationships, two complementary relationships, and two mixed or unresolved relationships.

Successive relationships were therefore the single most frequent pairwise class, and most remaining pairs were alternative rather than complementary. Together, these patterns indicate that representative taxa often occupied the same broader latent context without consistently occupying it simultaneously and frequently reached their strongest context contribution during different life stages. This provided the quantitative basis for the Successive descriptor. The term describes a population-level temporal offset across samples and developmental stages and does not imply an ordered replacement sequence within individual participants.

*Concentrated Bifidobacterium longum context*

The *B. longum* context was distinguished primarily by the concentration and life-stage consistency of its representative contributor set. The same five taxa occupied the five highest routed-mass-share positions across all five condensed life stages, yielding a representative-set union of five taxa, the minimum possible under the selection procedure.

The ten pairwise relationships among these taxa were heterogeneous: two were complementary, four alternative, three successive, and one mixed or unresolved. No single pairwise class therefore explained the architecture of this context. Instead, the Concentrated descriptor refers specifically to the small and repeatedly recovered contributor backbone across life stages. The steep head of the within-context taxon probability distribution was consistent with this concentration but was treated as supporting rather than defining evidence.

*Mixed Bacteroides vulgatus context*

The *B. vulgatus* context contained nine representative taxa and 36 pairwise relationships distributed across all four relationship classes. Eight pairs were complementary or complementary-leaning, 12 were alternative or alternative-leaning, 10 were successive or successive-leaning, and six were mixed or unresolved.

No relationship class accounted for a majority of evaluated pairs, and substantial support was distributed among complementary, alternative, and successive architectures. The Mixed descriptor therefore reflects the coexistence of several forms of internal organization within the same broader latent context rather than an absence of ecological structure.

*Overall architectural distinction among contexts*

Taken together, the four microbial contexts exhibited distinguishable forms of internal organization. The *P. copri* context had the greatest relative representation of complementary relationships, supporting a shared and aligned architecture. The *E. coli* context showed the largest number of successive relationships, supporting a predominantly staggered organization across life stages. The *B. longum* context was distinguished by the minimum-size, stage-consistent contributor set rather than by pairwise uniformity, while the *B. vulgatus* context showed the broadest distribution across pairwise relationship classes.

These quantitative patterns provide the post hoc basis for the descriptors Complementary, Successive, Concentrated, and Mixed, respectively. The descriptors summarize population-level organization within the learned microbial contexts and should not be interpreted as direct evidence of biochemical interaction, competition, or within-person ecological succession.

### Supplementary Results 2. Stochastic variability, representative-model stability and reproducibility of Discovery

#### *Across-configuration-space variability of LDA fits*

Independently initialized LDA fits produced different context compositions within the same configuration, even after optimal matching of their contexts. Thus, fixing the number of contexts and both filtering thresholds did not yield identical fitted models.

Separate principal coordinate analysis (PCoA) plots illustrated the context distributions recovered across random initializations and filtering settings within each K (Supplementary Fig. 6). The distributions occupied structured regions of composition space. The plots combined distinct contexts and multiple filtering settings, therefore they provided a descriptive overview of stochastic variability. Quantitative fit-to-fit comparisons within otherwise identical configurations showed the lowest and most concentrated matched PCoA-space distances at $K = 3 - 4$, with generally greater divergence from $K = 5$ onward, peaking at $K = 7$ before declining somewhat at higher $K$ values (Supplementary Table 8).

*Stability within the selected Discovery configuration*

The Within-selected-Discovery configuration analysis characterizes the stochastic variation addressed by the Discovery stabilization procedure and shows that the eligible realization set is concentrated around the representative solution. Principal coordinates analysis of the complete taxon-probability distributions showed that most context estimates occupied recurrent context-specific compositional regions, whereas Isolation Forest outliers were more dispersed and extended away from these dominant structures (Supplementary Fig. 7). The representative context estimates were positioned within the recurrent regions formed by the corresponding eligible realizations.

This pattern was also evident when compositional differences were quantified directly. Context estimates from eligible models generally showed low Jensen-Shannon distances from their corresponding representative contexts, whereas outlier models exhibited a substantially broader distribution of distances and contained the most divergent context estimates (Supplementary Fig. 7b).

Differences in context composition were reflected in downstream sample-taxon assignments. Eligible models showed consistently high exact assignment agreement with the representative realization, whereas outliers exhibited markedly greater variability and included realizations with substantially lower agreement (Supplementary Fig. 7c). Thus, stochastic differences among individual LDA fits were not restricted to small changes in context probability estimates but could, for atypical realizations, propagate to materially different individualized taxon-in-context assignments. Together, these results show that most LDA realizations generated under the selected Discovery configuration converged toward a recurrent context architecture, while a minority produced atypical solutions with greater compositional and assignment-level divergence.

*Reproducibility across independent Discovery trials*

The Between-Discovery trials analysis evaluates the stability of the entire Discovery workflow under varying independent runs of the entire workflow. Across independent Discovery runs, all Discovery trials selected four microbial contexts on CMD data, although filtering thresholds and the number of retained taxa varied across runs (Supplementary Table 2). After matching microbial contexts by taxonomic composition, the largest within-context Jensen-Shannon distance was 0.18, below the smallest between-context distance of 0.87 (Supplementary Fig. 3a,b). Assignment-level reproducibility was similarly high: 98% of sample-taxon observations with valid assignments in all runs showed unanimous agreement, with context-specific unanimous agreement ranging from 99% to 100%; pairwise exact assignment agreement across runs was consistently above 98% (Supplementary Table 3).

### Supplementary Results 3. Correspondence of microbial contexts across datasets and taxonomic resolutions

*Correspondence between CMD contexts and contexts recovered from the combined genus-level dataset*

Discovery identified four species-level microbial contexts in CMD reference dataset and five genus-level microbial contexts in the larger GMR-Compendium dataset, which integrated shotgun metagenomic and 16S profiles (Supplementary Fig. 4a). We compared their taxonomic composition and age-associated activation patterns to assess correspondence between the two analyses.

Several lineage-level correspondences were observed. The CMD Mixed *B. vulgatus* context showed compositional similarity with  the GMR-Compendium *Bacteroides*-interacting context, both dominated by *Bacteroides*-associated gut anaerobes. The CMD Successive *E. coli* context showed broad correspondence with the GMR-Compendium *Citrobacter*-interacting context, reflecting an *Enterobacteriaceae*-associated configuration. The CMD Concentrated *B. longum* context showed similarity to the GMR-Compendium Streptococcus-interacting context, which included early-life-associated genera such as *Streptococcus* and *Bifidobacterium*. The CMD Complementary *P. copri* context corresponded more broadly to two adult-associated genus-level contexts in GMR-Compendium: *Prevotella*-interacting and *Agathobacter*-interacting.

Age-associated activation patterns were also broadly consistent across datasets (Fig. 1d; Supplementary Fig. 4b). Early-life and Enterobacteriaceae-associated contexts showed higher activation early in life and declined with age, whereas adult-associated anaerobic contexts became more prominent later in life. These results support broad correspondence in microbial context composition and age-associated patterns between CMD and the combined genus-level dataset.

### Supplementary Results 4. Conceptual benchmarking against enterosignatures

#### *Discovery contexts show broad compositional concordance with enterosignatures*

To determine whether Discovery recovered previously described population-level microbiome structures, we compared microbial contexts with enterosignatures[4] using Jensen-Shannon distance. Four of the five microbial contexts (80%) showed close correspondence to a single enterosignature, with distances ranging from 0.16 to 0.33 (median, 0.23). The closest matches were the Bacteroides context with ES-Bact (0.16), the Escherichia context with ES-Esch (0.21), the Agathobacter context with ES-Firm (0.25) and the Prevotella context with ES-Prev (0.33).

These four context-enterosignature pairs shared the same leading taxon and broadly similar high-contribution taxa, although their relative contributions differed. This concordance supports the interpretation that Discovery recovers recognizable high-level microbial configurations from complex microbiome data. The comparison also showed that microbial contexts and enterosignatures differ in how they weight taxa within these configurations.

Notably, the two frameworks are closely related in construction: non-negative matrix factorization under a Kullback-Leibler objective is equivalent to probabilistic latent semantic analysis[43], of which latent Dirichlet allocation[5] is the Bayesian extension, and enterosignatures are fitted with this Kullback-Leibler objective[4]. Differences between paired components therefore reflect the priors and the particular fitted solution rather than a different treatment of taxon abundance. This benchmark is accordingly interpreted as convergent validation of the context reference space at the population level. What Discovery adds is a per-sample, taxon-level context assignment.

**Supplementary Results 5. Relationship between context-responsive taxa and context pivots**

Of the 64 taxa identified as context-responsive in the abundance-controlled analysis, 37 also satisfied the context-pivot criterion after adjustment for the sample-level context activation profile and calibration against composition-compatible null assignments (Supplementary Methods 5). Twenty-seven taxa were supported only by the abundance-controlled analysis.

For 23 of these 27 taxa, the assignment effect was substantially attenuated after inclusion of the sample-level context activation profile. This group included *Escherichia coli*, *Collinsella aerofaciens* and *Bifidobacterium adolescentis*. Four additional taxa retained a significant composition-controlled nested effect but did not exceed the composition-compatible null. Conversely, five context pivots were not detected by the abundance-controlled analysis but met the context-pivot criterion after community adjustment. Thus, the two analyses identified overlapping but non-identical sets of taxa, with 37 taxa supported under both analytical criteria.

**Supplementary Results 6. Additional evaluation and sensitivity analyses of recurrent UTI prediction**

*Temporal characteristics of the two-visit recurrence analysis*

The two retained ordinal samples were concentrated at the earliest scheduled study timepoints (Supplementary Table 4). Among the 105 participants included in the recurrence analysis, 95 of 105 first observations (90.5%) corresponded to study timepoint 1, while nine (8.6%) corresponded to timepoint 2 and one (1.0%) to timepoint 4. The first observation occurred at a median collection day of 0 (range 0-14 days).

The second observed sample most commonly corresponded to study timepoint 2 (73/105, 69.5%). A further 16 participants (15.2%) contributed their second observation at timepoint 3, such that 84.8% of second observations occurred by timepoint 3. Eleven second observations (10.5%) occurred at timepoint 4 and only five (4.8%) occurred at timepoints 5-6. The median collection day for the second observation was 10 days (range 2-40 days).

The temporal distribution was similar between outcome groups. For both recurrent and non-recurrent participants, the median collection day was 0 days for the first observation and 10 days for the second observation. At the first ordinal visit, 90.5% of non-recurrent and 90.5% of recurrent participants were sampled at study timepoint 1. At the second ordinal visit, 68.3% of non-recurrent and 71.4% of recurrent participants were sampled at study timepoint 2. Thus, there was no obvious gross shift in the timing of the retained observations between recurrence groups.

Timing-only models nevertheless showed weak residual outcome separation. Across the nine combinations of timing representation and probe model, direction-free AUROC estimates ranged from approximately 0.51 to 0.58. The largest direction-free AUROC was 0.575 for the ordinal-timepoint linear logistic probe. None of the timing probes satisfied the pre-specified practical-equivalence criterion because their one-sided upper confidence bounds exceeded 0.57. Thus, timing alone showed substantially weaker discrimination than the microbiome context model, but the available data were insufficient to demonstrate that timing-related information was completely negligible.

Together, these results reduce concern that the recurrence signal was driven predominantly by the sampling calendar. However, exact recurrence dates were unavailable, therefore temporal precedence of both retained samples could not be independently verified. The analysis should therefore be interpreted as internally cross-validated prediction from early two-sample microbiome profiles rather than as prospectively landmarked pre-recurrence prediction.

*Comparison with thresholded presence and feature-space separation*

Supervised sample-level projections of abundance and context-state features for the same 24 selected taxa showed greater descriptive separation between samples from recurrent and non-recurrent participants for the context representation. The Euclidean distance between outcome-group centroids, calculated after standardizing both projected axes to unit standard deviation, was 0.751 for standardized raw abundance and 1.164 for context states (Fig. 6c,d). Substantial overlap remained between the groups, and these projections were used only to visualize the corresponding feature spaces rather than as an independent assessment of predictive performance.

Because taxon-in-context assignment required taxa to meet the 0.1% abundance floor, a threshold-matched abundance-presence representation was evaluated as an additional comparator. The abundance-presence model achieved an AUROC of 0.536 (95% CI, 0.415 to 0.661), compared with 0.707 (95% CI, 0.600 to 0.805) for the context-only model. The estimated difference between context and threshold-matched presence was 0.172 AUROC (95% CI, 0.060 to 0.287). Comprehensive performance across the matched microbiome representations is provided in Supplementary Table 5.

*Robustness of recurrence discrimination across linear classifiers*

In a separate sensitivity analysis using a common 226-taxon universe, the relative performance of the three microbiome representations was evaluated across five linear classifiers and 20 repetitions of nested cross-validation, with AUROC calculated from participant-level scores averaged across repetitions (Supplementary Table 6). Taxon-in-context AUROC ranged from 0.653 to 0.706 across classifiers, compared with 0.454 to 0.487 for CLR abundance and 0.520 to 0.570 for thresholded presence. Paired context-minus-abundance differences ranged from 0.179 to 0.231 and context-minus-presence differences from 0.110 to 0.164; all corresponding conditional unadjusted 95% bootstrap intervals excluded zero. Taxon-in-context exceeded CLR abundance in 19–20 of 20 repetitions and exceeded thresholded presence in 18–20 of 20 repetitions, depending on classifier. These results indicate that the representation-level performance advantage was generally consistent across the tested classifiers within this cohort.

Complementary label-permutation analyses, using a five-repetition observed reference, were directionally consistent but yielded weaker multiplicity-adjusted evidence. After Holm correction, the L1 logistic-regression comparison against CLR abundance remained significant ( $P_{adj} = 0.040$); the remaining abundance comparisons had adjusted $P_{adj} = 0.072$, and all thresholded-presence comparisons had adjusted $P_{adj} = 0.090$. These tests evaluate a global absence-of-association null and therefore complement rather than duplicate the paired performance comparisons.

<u>*Stable taxon-in-context predictors*</u>

Fifteen taxon-in-context-visit features met the pre-specified stability criterion in the primary context-only model, comprising ten features positively associated with recurrence and five negatively associated with recurrence (Fig. 6e; Supplementary Table 7). Among the largest positive coefficients were *Eubacterium eligens* assigned to the Mixed *B. vulgatus* context at visit 1 and *Streptococcus parasanguinis* assigned to the Complementary *P. copri* context at visit 2. Negative coefficients included *Hungatella hathewayi*, *Clostridium innocuum* and *Alistipes finegoldii* assigned to the Mixed *B. vulgatus* context. Features assigned to the Mixed *B. vulgatus* context occurred in both coefficient directions.

Supplementary Discussion

### **Biological interpretation and future directions**

Beyond these context level summaries, taxon-in-context analysis revealed that the same bacterial species can inhabit different microbial contexts across individual microbiomes. For example, *Bifidobacterium adolescentis* occupied either the Concentrated *B. longum* or the Complementary *P. copri* context across the lifespan. In our analysis, *B. adolescentis* occupied the Concentrated *B. longum* context from early through later life; co-occurrence of *B. adolescentis* and *B. longum* in metagenomic datasets has been associated with healthy individuals[44]. In the *P. copri* context, *B. adolescentis* appears alongside butyrate associated taxa, mainly during adulthood, consistent with reported positive associations between bifidobacteria and butyrate producing organisms in healthy individuals[44]. *Bifidobacterium adolescentis* is a recognized utilizer of plant derived carbohydrates and may support butyrate production through cross feeding with butyrate producing bacteria[15]. Similarly, *F. prausnitzii* occupied either the Complementary *P. copri* context or the Mixed *B. vulgatus* context, consistent with its ability to use fermentation products generated by primary degraders while contributing to shared metabolic outputs[45,46]. *Ruminococcus gnavus* is another multicontext taxon, occupying either the Successive *E. coli* or the Mixed *B. vulgatus* context across the lifespan. Its assignments were distributed between the two contexts during childhood but became concentrated in the Mixed *B. vulgatus* context in adulthood, with a minority remaining in the Successive *E. coli* context. *Ruminococcus gnavus* has been associated with diseases in early and later life; resolving its ecological context may help distinguish the community settings in which those associations arise[47–52].

The lifespan structure of these assignments revealed variation in microbial context organization across age groups. Early life showed the highest ecological context flexibility, consistent with the rapid establishment of the gut microbiome and the successive colonization of microbes adapted to human milk oligosaccharides, followed by microbes involved in the fermentation of fibers and phytochemicals once weaning occurs[16]. In the early life dataset, 22 taxa showed delivery mode associated differences in the contexts they occupied after accounting for abundance. Across delivery groups, these microbes were preferentially embedded in the Mixed *B. vulgatus* context in vaginally delivered infants and the Successive *E. coli* context in infants delivered by C section. This redistribution was consistent with the delayed colonization by *Bacteroides* species reported after C section delivery[53–55], with reduced occupancy of the Mixed *B. vulgatus* context and greater occupancy of the Successive *E. coli* context. The Mixed *B. vulgatus* context includes *Bacteroides* species previously associated with vaginal delivery and breastfeeding[56], whereas the Successive *E. coli* context is enriched in microbes with opportunistic potential that have been reported after C section delivery[14]. These observations motivate testing whether the functional contribution of these microbes differs between ecological contexts associated with delivery mode. For example, *Ruminococcus gnavus* has been associated with allergy risk in early life[47–49]. Whether that association differs when the species inhabits the Mixed *B. vulgatus* or Successive *E. coli* context is a specific, testable hypothesis.

The gradual decline in context entropy across adult age groups in CMD indicates more consistent context occupancy for particular species across people of similar age. Previous multiomics and microbiome studies have reported inflection points around mid adulthood and later life, including transitions around 40 and 60 years of age[57]. Those studies suggest non linear aging microbiome trajectories[57,58]. Our findings add a complementary view by showing that the ecological contexts occupied by microbes also differ across adult age groups. Longitudinal sampling is needed to determine how these patterns develop within an individual.

To explore whether taxon-in-context organization distinguished contrasting aging associated microbiome configurations, we stratified samples using elevated activation of the Successive *E. coli* context outside early life. Because this context is enriched in facultative anaerobes/aerobes and microbes with opportunistic potential, the terms typical and atypical are used descriptively and do not denote independently validated health states. CMD was restricted to samples annotated as healthy after exclusions based on available metadata; typical and atypical therefore describe relative configurations within this reference collection. As expected, the *E. coli* context enriched group was most evident in later life and showed higher activation of the *E. coli* context together with lower activation of the *B. vulgatus* and *P. copri* contexts. The broader pattern resembles gut microbial signatures previously associated with dysbiosis, inflammation and age related disease[13,50–52,59–64], but independent clinical phenotypes will be needed to establish its health relevance.

Taxon-in-context features made explicit associations not captured by focal taxon relative abundance alone. The context a microbe occupies provides a compact, per sample readout of the broader community configuration, so that even a single species can index whether an individual exhibits the typical or atypical configuration. *Streptococcus parasanguinis* occupied either the Successive *E. coli* context, characteristic of the atypical configuration, or the Complementary *P. copri* context, characteristic of the typical configuration, reflecting the community state in which the species was embedded rather than a change in its abundance. A similar pattern was observed for *Ruminococcus gnavus*. These alternative ecological embeddings do not establish different functions, but they generate a testable hypothesis that occupation of the Successive *E. coli* context is associated with a community enriched in gut and oral microbes with opportunistic potential. Both species have been reported in age related disease cohorts. Resolving them in context may help determine whether those associations depend on ecological setting and whether their context specific occurrence has value as an aging biomarker[50–52]. Together, these examples show that the same species can index different aging associated configurations depending on its assigned context, rather than carrying a single context independent interpretation.

Context responsive taxa and context pivots provide complementary views of age associated organization. Context pivots, such as *Faecalibacterium prausnitzii*, improved age model fit after accounting for focal abundance and additive context activation effects, and exceeded the specified composition compatible null. The two groups make complementary points. Context responsive taxa show that a single taxon assignment can index the configuration of the whole community, a compression not explicit in static representations such as enterosignatures. Context pivots identify taxon and context pairings whose age associations were not captured by the focal abundance and additive context activation terms in the fitted models. These pairings provide candidates for investigating ecological participation, without establishing information beyond the full community abundance profile. Context pivots should not be interpreted as microbes that necessarily initiate or maintain a microbial context: whether some pivots precede or facilitate broader recruitment into a context is an important assembly hypothesis that will require extensive longitudinal data or experimental testing.

Predictive features represented a mixture of recurrence and non recurrence associated microbial configurations (Supplementary Table 7). Assignment of specific gut commensals to the Mixed *B. vulgatus* context at the first two observed samples was associated with non recurrence. Specifically, assignment of *Bacteroides xylanisolvens*, *Clostridium innocuum*, *Alistipes finegoldii*, and *Hungatella hathewayi* to the *B. vulgatus* context was consistently associated with non recurrence. In contrast, recurrence was associated with, among others, *Eubacterium eligens* and *Parabacteroides goldsteinii* occupying the *B. vulgatus* context, and *Streptococcus parasanguinis* and *Ruminococcaceae bacterium D5* occupying the *P. copri* context.

Together, these findings support a context aware view of the human gut microbiome. Microbial contexts provide a reproducible representation of latent taxonomic configurations inferred from whole community structure, but the main advance is the ability to resolve the ecological setting inhabited by each microbe within an individual microbiome and to follow changes in that setting over time. This makes it possible to distinguish microbes anchored to one context from those that inhabit different contexts across microbiomes, to detect age associated reorganizations of microbiome ecology and to test whether taxon-in-context features improve prediction of clinical outcomes or intervention responses beyond matched abundance representations. The resulting framework suggests that precision microbiome analysis should not only ask which microbes are present, but also which contexts they occupy, how taxon-in-context organization changes across life and perturbation, and whether context shifts can be modulated by diet or other interventions.

Strain resolved context assignment is a particularly important next step. Closely related strains can differ in metabolic capacity and host interaction[17,18]; probiotic and other biotic studies could test whether response depends on the context in which an administered or resident strain becomes embedded. Whether context flexibility is greater at strain than species level remains a hypothesis requiring direct strain resolved validation.

More generally, context assignment also reframes how interventions might be conceived in terms of shifting the ecological settings in which microbes are embedded rather than only raising or lowering their abundance, a possibility that remains to be tested directly. In intervention studies, baseline context activation and taxon-in-context assignment profiles could be evaluated alongside baseline diet and host biology as predictors of response, while longitudinal sampling could test whether intervention efficacy is accompanied by specific context shifts even when abundance changes are modest. This provides a foundation for context aware precision nutrition and for future studies aimed at understanding how individual microbiome configurations shape host health[65–68].

Supplementary Figures

### Supplementary Fig. 1: Stability-breakpoint selection of the number of microbial contexts

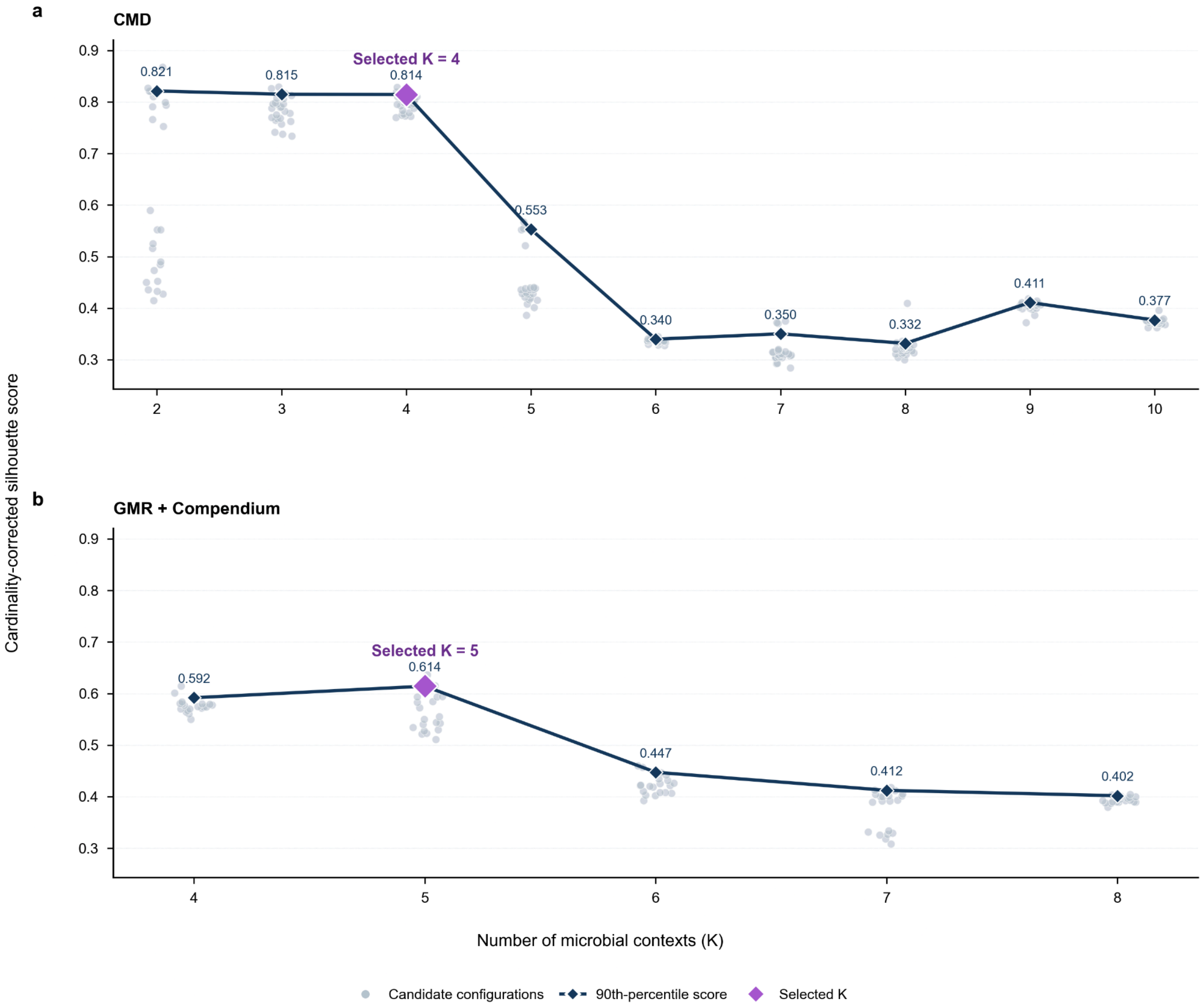


**Supplementary Fig. 1: Stability-breakpoint selection of the number of microbial contexts. a,** Cardinality-corrected silhouette scores across candidate context numbers in CMD. **b,** Corresponding scores in the GMR-Compendium dataset. Light points represent individual abundance and prevalence threshold configurations; connected symbols represent the 90th percentile of the configuration scores at each $K$; and, highlighted symbol indicates the selected $K$. CMD showed a stable $K = 3 - 4$ pair followed by a persistent decline beginning at $K = 5$, supporting $K = 4$. The selected abundance and prevalence thresholds are 0.003 and 0.005, respectively, resulting in 275 retained species. GMR-Compendium showed a stable $K = 4 - 5$ pair followed by a persistent decline beginning at $K = 6$, supporting $K = 5$. The selected abundance and prevalence thresholds are 0.005 and 0.008, respectively, resulting in 74 retained genera.

### Supplementary Fig. 2: Taxon contribution patterns and pairwise metrics underlying microbial context descriptors

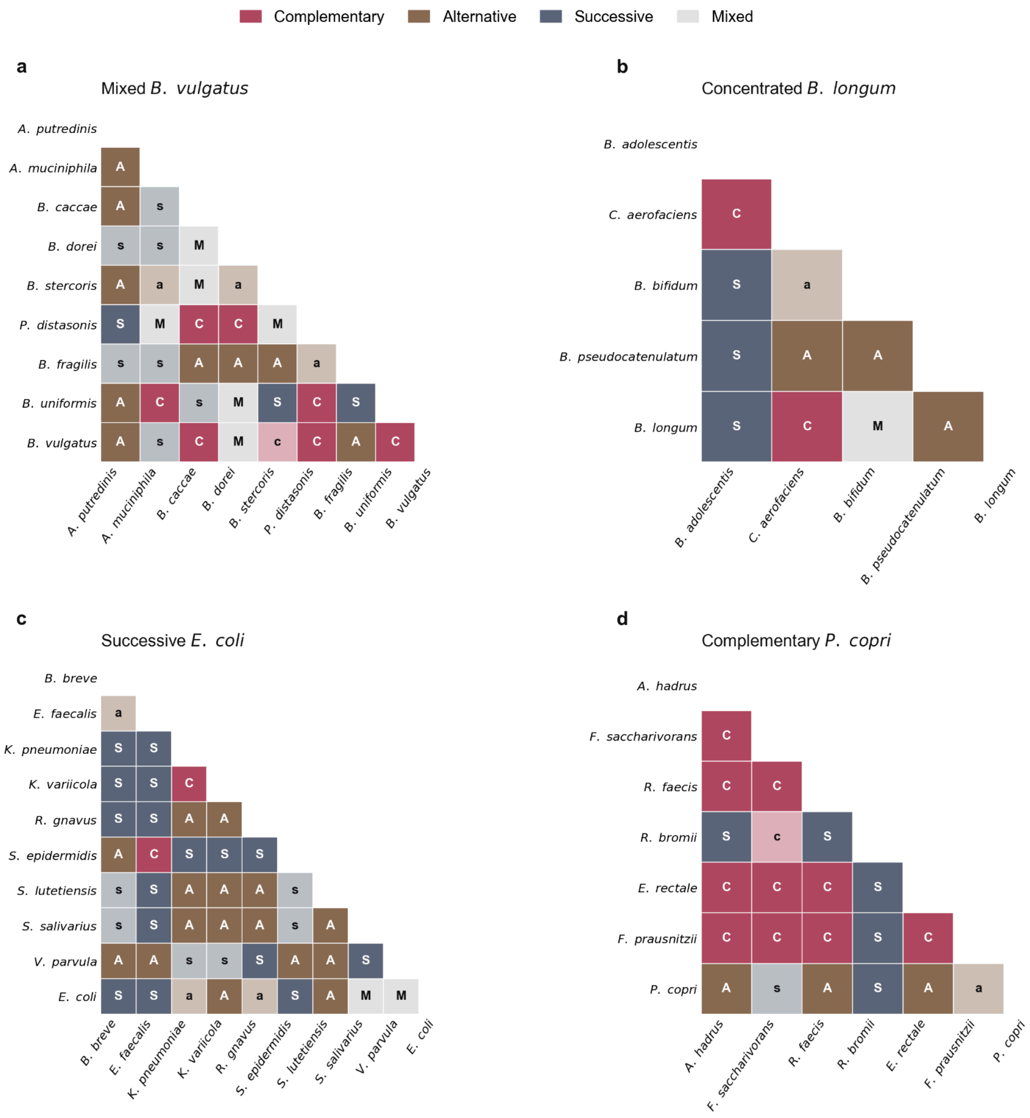


**Supplementary Fig. 2: Taxon contribution patterns and pairwise metrics underlying microbial context descriptors.** Pairwise temporal relationships among member taxa were classified as complementary (C), alternative (A), successive (S) or mixed (M) and displayed as lower-triangular matrices for the **a,** Mixed B. vulgatus, **b,** Concentrated B. longum, **c,** Successive E. coli and **d,** Complementary P. copri contexts. Each cell represents the relationship between a taxon pair, with colours and letter codes indicating the inferred relationship class. The distinct distributions of pairwise relationships provide the basis for the ecological characterization of each microbial context.

### Supplementary Fig. 3: Between-Discovery trials analysis identifies compositionally distinct microbial contexts across independent trials

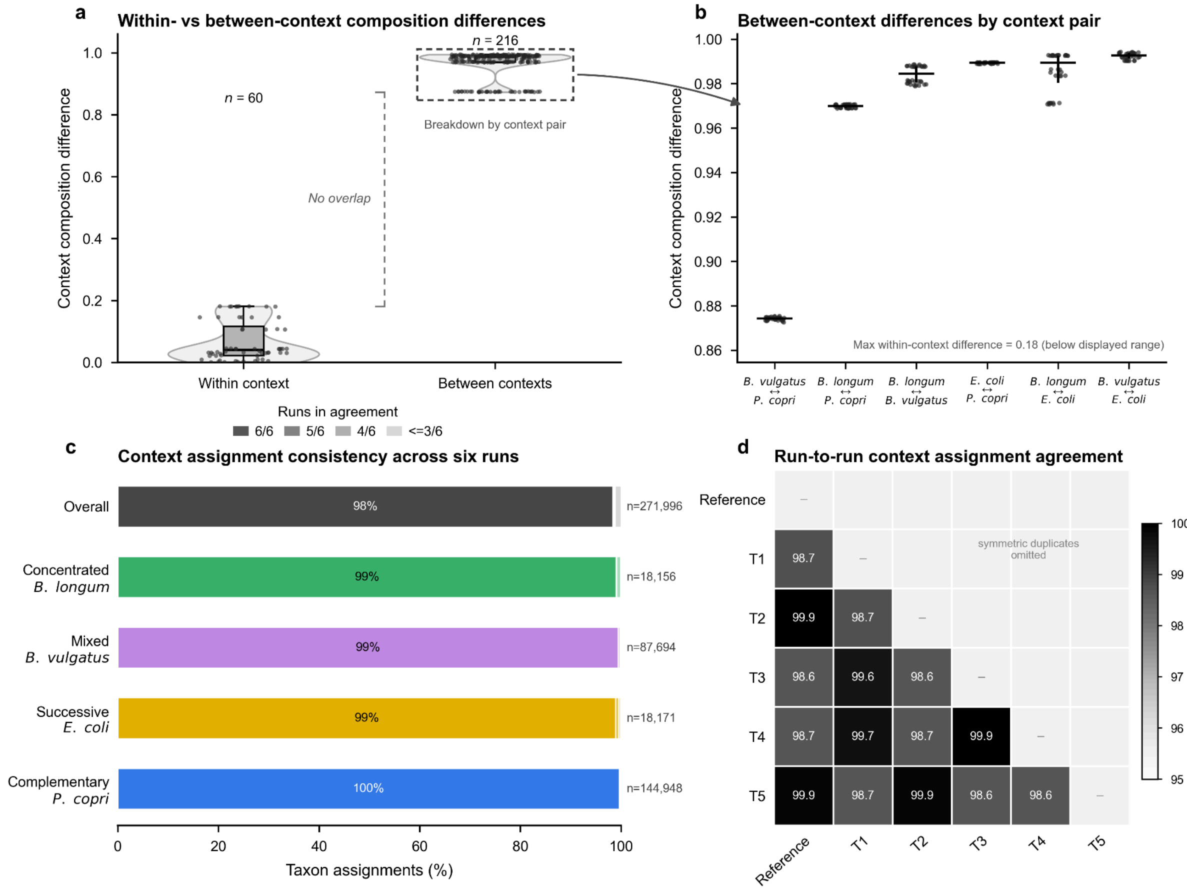


**Supplementary Fig. 3: Between-Discovery trials analysis identifies compositionally distinct microbial contexts across independent trials.** Six Discovery runs from independent random-seed batches were aligned to a reference architecture using one-to-one matching of context compositions based on Jensen-Shannon distance (JSD). **a,** Within-context compositional differences were consistently smaller than between-context differences, with no overlap between the two distributions. **b,** Between-context JSDs shown separately for each context pair. **c,** Agreement of sample-taxon context assignments across all six runs, shown overall and by aligned microbial context. **d,** Pairwise exact assignment agreement between runs. Context alignment was determined from composition alone and then applied to the assignment data. Across trials, Discovery recovered compositionally distinct but reproducibly aligned contexts and highly consistent taxon assignments, demonstrating robustness to independent stochastic initialization.

### Supplementary Fig. 4: Microbial contexts and their corresponding developmental age-related patterns based on GMR-Compendium dataset

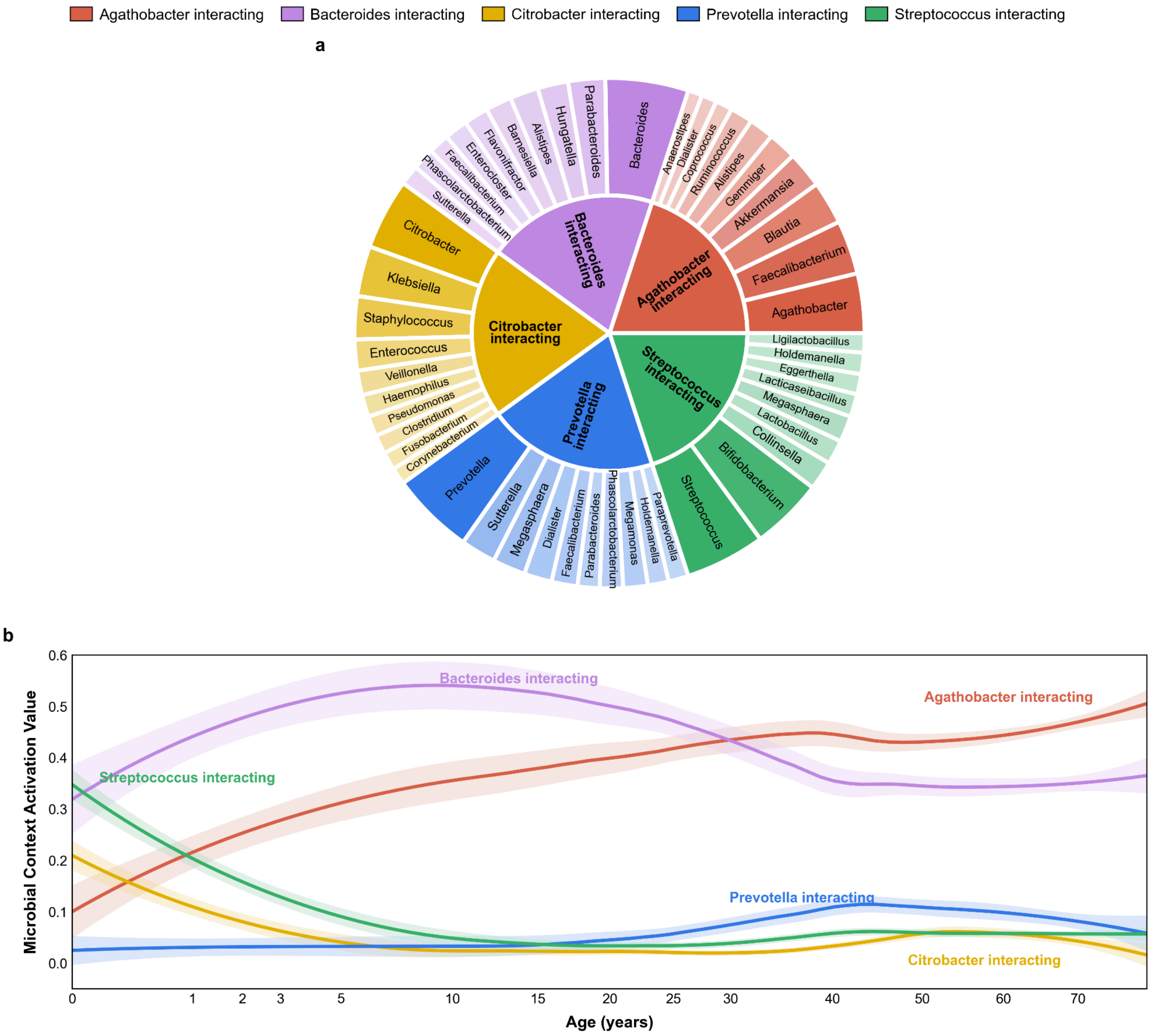


**Supplementary Fig. 4: Microbial contexts and corresponding age-associated patterns in the GMR-Compendium dataset. a,** Sunburst representation of the five inferred microbial contexts. Inner sectors denote contexts and outer sectors show the ten highest-weighted genera within each context, with wedge size and shading reflecting context-composition weights used for visualization. **b,** Age-associated context activation profiles. Activation values were normalized across contexts. Samples with age −1, zero abundance across all supplied genera or uniform normalized activation across contexts were excluded before age ordering, window averaging and smoothing. Retained activation profiles were averaged within consecutive 50-sample age-ordered windows and smoothed on a signed-square-root age scale; shaded bands indicate 95% confidence intervals.

### Supplementary Fig. 5: Ecological context flexibility based on GMR-Compendium dataset

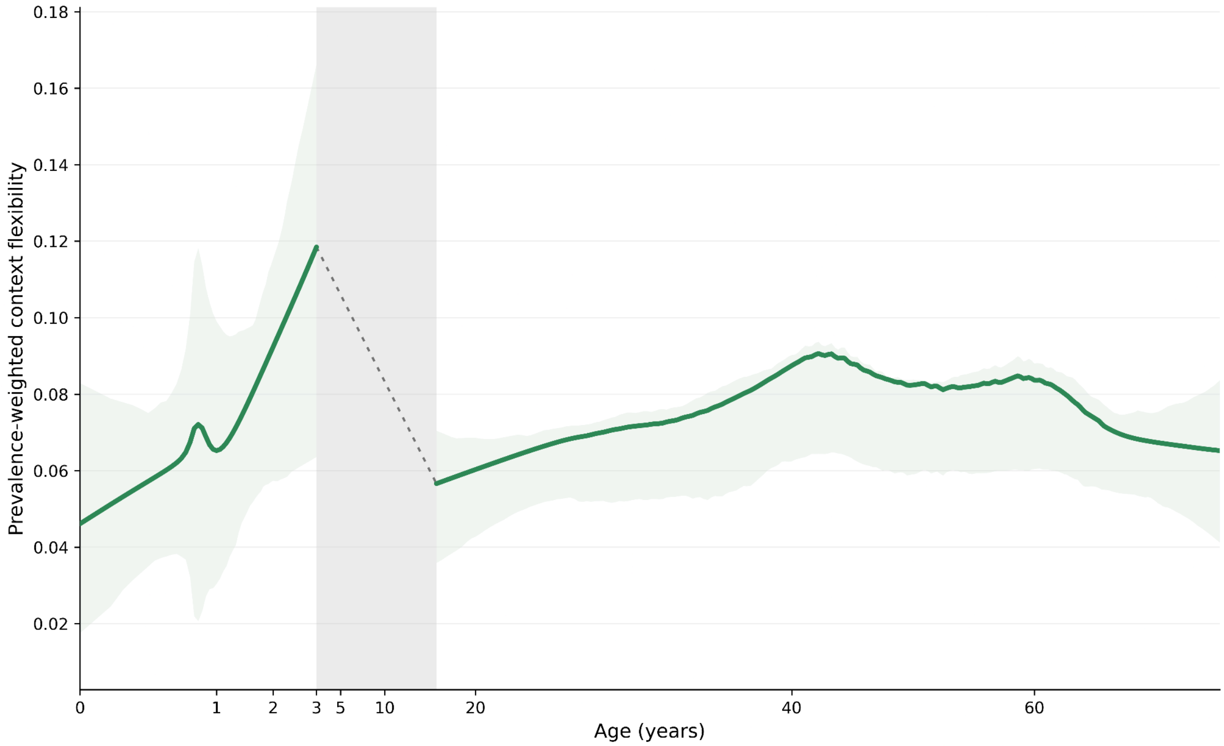


**Supplementary Fig. 5: Ecological context flexibility based on GMR-Compendium dataset.** Prevalence-weighted context flexibility was calculated from genus-level context assignments and summarized across age using study-balanced resampling. Supported early-life (0-3 years) and adult (≥18 years) segments were fitted independently using LOWESS; shaded bands indicate 95% bootstrap intervals. The grey region denotes the lower-support interval, with the dashed line shown only as an endpoint connector and not as an estimated trajectory. The GMR+Compendium analysis reproduced the CMD pattern of elevated ecological flexibility in early childhood followed by lower flexibility in adulthood, despite differences in taxonomic resolution and sequencing platform.

### Supplementary Fig. 6: Context composition across LDA fits and filtering configurations

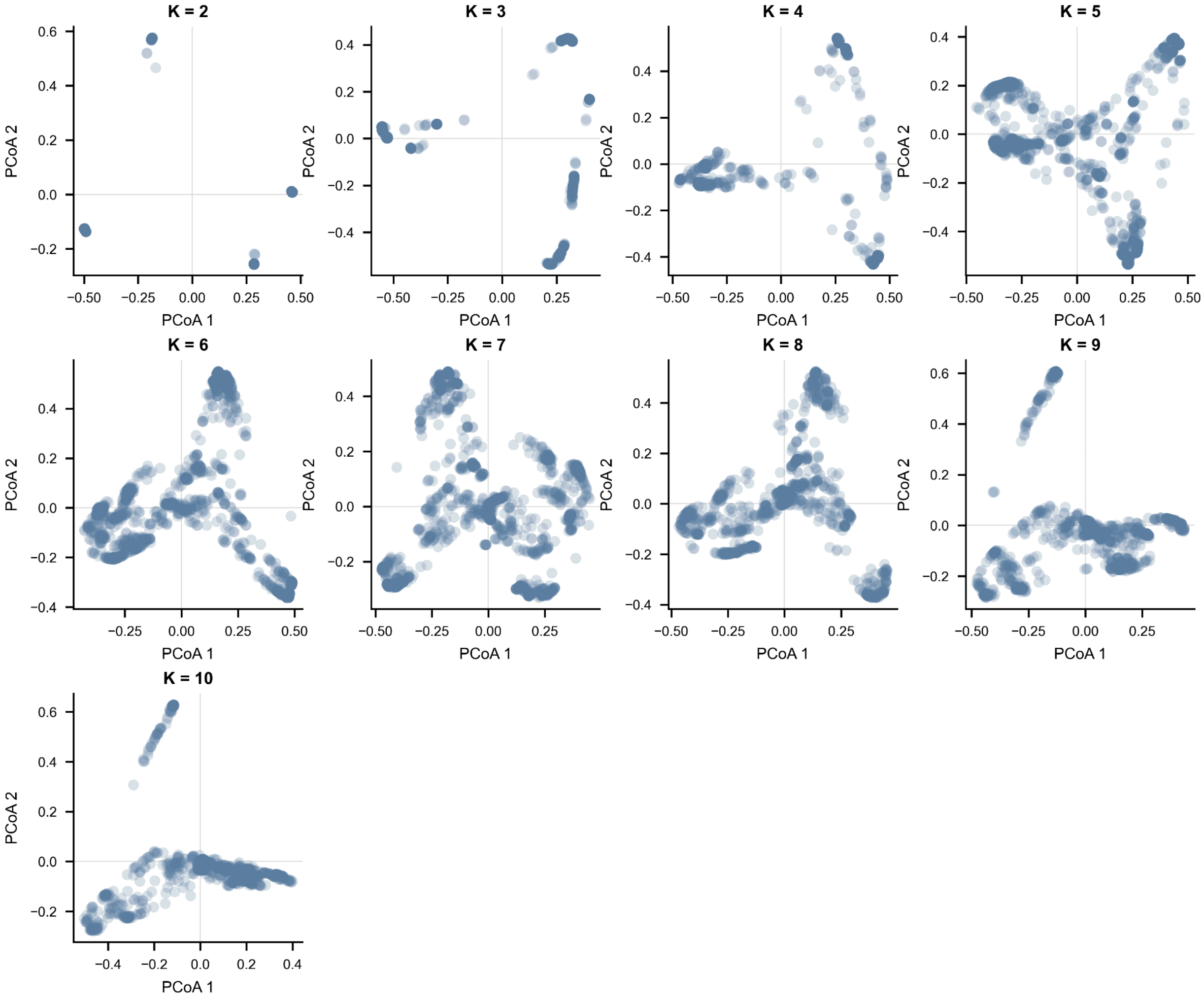


**Supplementary Fig. 6: Context composition across LDA fits and filtering configurations.** Principal coordinate analysis (PCoA) of inferred microbial context compositions across candidate numbers of contexts $K = 2 - 10$. For each $K$, context-composition vectors from the evaluated filtering configurations were pooled, pairwise Jensen-Shannon distances were calculated, and PCoA was performed independently. Each point represents one inferred microbial context from a sampled LDA realization. The dispersion of points within each panel illustrates the variability in context composition arising across filtering configurations and model realizations for a fixed $K$, while the clustering of points indicates recurring context-composition patterns within each $K$. Because ordination was performed separately for each $K$, axis orientation and absolute coordinates should be interpreted only within each panel and not compared directly across $K$.

### Supplementary Fig. 7: Within-Discovery configuration analysis reveals a recurrent context structure and stable downstream assignments

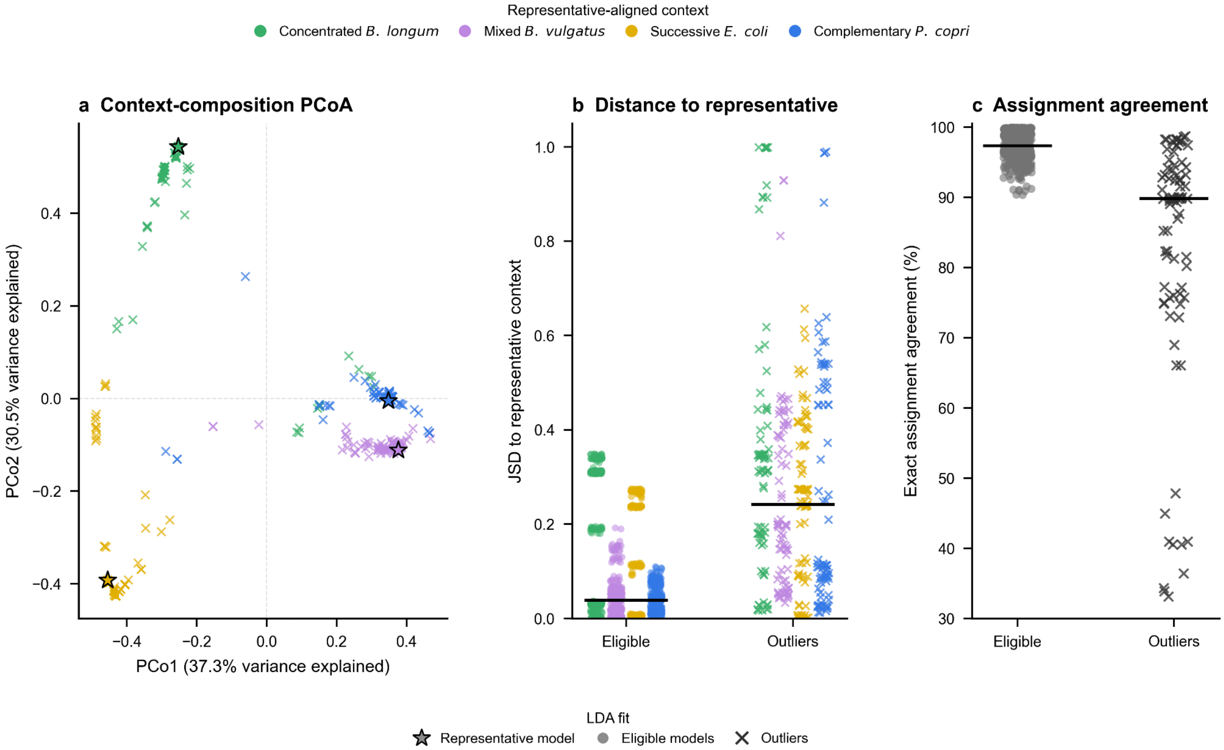


**Supplementary Fig. 7: Within-Discovery configuration analysis reveals a recurrent context structure and stable downstream assignments. a,** PCoA of context compositions from complete LDA fits based on pairwise Jensen-Shannon distances (JSDs). Contexts were aligned to the selected representative model; colours denote representative-aligned contexts, circles denote eligible fits, crosses denote Isolation Forest outliers and stars denote the representative model. **b,** JSD of each aligned context to its corresponding representative context. **c,** Exact sample-taxon context-assignment agreement with the representative model. Black horizontal lines indicate medians. Eligible fits showed recurrent context compositions, with lower compositional distances and consistently high assignment agreement, whereas outlier fits exhibited greater compositional and assignment variability.

Supplementary Tables

### Supplementary Table 1. Definitions of key terms used in the taxon-in-context framework

| Term | Definition |
|---|---|
| *Microbial context* | A latent taxonomic configuration learned from whole-community composition across samples. Each context is represented by a probability distribution over taxa and provides a shared reference for interpreting individual microbiomes. |
| *Context activation* | The estimated contribution of a microbial context to a sample's overall context mixture. Activation values sum to one across contexts within each sample. |
| *Taxon-in-context* | A representation of a taxon together with its inferred microbial context in an individual microbiome, making contextual information implicit in community composition explicit at taxon and sample resolution. |
| *Context occupancy* | The inferred context associated with a taxon in a particular sample. In this study, occupancy is determined by the context with the highest posterior probability for that taxon-sample observation. |
| *Taxon-in-context profile* | The collection of taxon-specific context occupancies within a microbiome, describing which contexts individual microbes occupy. |
| *Single-context taxon* | A taxon observed in only one inferred microbial context across the analysed samples. This designation applies to the dataset and model examined and does not imply an intrinsically fixed ecological identity. |
| *Multicontext taxon* | A taxon observed in more than one inferred microbial context across the analysed samples. This designation describes variation across samples and does not, by itself, establish switching within an individual. |
| *Ecological context flexibility* | The extent to which a taxon occupies different inferred microbial contexts across individual microbiomes. In the lifespan analysis, flexibility describes variation among individuals within age groups. |

| | |
|---|---|
| *Context entropy* | Normalized Shannon entropy of a taxon's context occupancy distribution across samples. Higher values indicate occupancy distributed more evenly across contexts; lower values indicate concentration in fewer contexts. Community-level estimates combine taxon-level values using prevalence weights. |
| *Ecological individuality* | Between-person variation in inferred taxon-in-context organization, through which the same taxon can occupy different microbial configurations in different individuals. |
| *Within-person context switching* | An observed change in the inferred context occupied by the same taxon across longitudinal samples from one individual. Strict switches occur between learned contexts and exclude transitions involving the no-context state. |
| *Strict context switch* | A strict context switch is a change between two different learned contexts in consecutive samples and excludes transitions involving other contexts or the no-context-assignment state. |
| *Context-responsive taxon* | A taxon whose context occupancy is associated with a host variable after accounting for its abundance. In the age analysis, context-responsive taxa improve age model fit relative to a flexible focal-abundance model. |
| *Context pivot* | An operational category from the age analysis: a taxon whose context occupancy improves model fit after adjustment for focal abundance and additive context activation effects, with an improvement exceeding the specified context-weight-based null. |
| *No-context state* | A retained taxon-sample observation for which no context is resolved under the preprocessing and inference rules. This is not an additional microbial context and does not imply ecological isolation. |
| *Discovery* | The ensemble-based LDA workflow developed in this study to select a reproducible context reference model and infer context activation and taxon-in-context profiles. |
| *Discovery Configuration* | A candidate combination of the number of microbial contexts (K), the minimum relative-abundance threshold and the sample-prevalence threshold. Candidate configurations are compared using the cardinality-corrected silhouette score. |

### **Supplementary Table 2. Selected Discovery configurations across the reference run and five reproducibility trials**

| Run | Number of contexts, K | Abundance threshold | Prevalence threshold | Penalized silhouette score | Retained taxa, n |
|---|---|---|---|---|---|
| **Reference run** | 4 | 0.003 | 0.005 | 0.828 | 275 |
| Trial 1 | 4 | 0.003 | 0.003 | 0.815 | 314 |
| Trial 2 | 4 | 0.003 | 0.005 | 0.820 | 275 |
| Trial 3 | 4 | 0.001 | 0.003 | 0.817 | 387 |
| Trial 4 | 4 | 0.001 | 0.003 | 0.812 | 387 |
| Trial 5 | 4 | 0.005 | 0.004 | 0.809 | 261 |

*Note.* The reference run corresponds to the original Discovery run used for manuscript analyses. Original runs 2-6 are relabeled here as Trials 1-5. Abundance and prevalence thresholds are shown as proportions. Cardinality-corrected silhouette scores are rounded to three decimal places.

### **Supplementary Table 3. Assignment-level reproducibility across six independent Discovery runs**

Primary analyses were restricted to sample-taxon observations with a valid context assignment in all six runs. Context identities were fixed using context matching based on taxonomic composition before assignment reproducibility was evaluated. The sensitivity analysis used the union of sample-taxon observations and treated observations absent from a run as No assignment.

#### a. Overall assignment reproducibility

| Measure | Primary analysis: shared taxon-sample observations | Sensitivity analysis: missing context assignments |
|---|---|---|

| | | |
|---|---|---|
| Analysis denominator | 271,996 common observations | 279,964 union observations |
| Common-assignment coverage | 271,996/279,964 (97.15%) | Not applicable |
| Unanimous 6/6 consensus | 267,655/271,996 (98.40%) | 95.60% |
| Strong consensus (>=5/6) | 268,282/271,996 (98.63%) | 96.85% |
| Pairwise exact agreement | Median 98.69%; range 98.62%-99.92% | Median 97.87% |

**b. Consensus stability by reference-aligned context**

| Consensus context | n | 6/6 | 5/6 | 4/6 | <=3/6 |
|---|---|---|---|---|---|
| **Overall** | **271,996** | **267,655 (98.40%)** | **627 (0.23%)** | **666 (0.24%)** | **3,048 (1.12%)** |
| Concentrated *B. longum* | 18,156 | 17,980 (99.03%) | 23 (0.13%) | 144 (0.79%) | 9 (0.05%) |
| Mixed *B. vulgatus* | 87,694 | 87,224 (99.46%) | 264 (0.30%) | 200 (0.23%) | 6 (<0.01%) |
| Successive *E. coli* | 18,171 | 17,976 (98.93%) | 111 (0.61%) | 80 (0.44%) | 4 (0.02%) |
| Complementary *P. copri* | 144,948 | 144,475 (99.67%) | 229 (0.16%) | 242 (0.17%) | 2 (<0.01%) |

| | | | | | |
|---|---|---|---|---|---|
| Tie (no unique consensus context) | 3,027 | 0 (0.00%) | 0 (0.00%) | 0 (0.00%) | 3,027 (100.00%) |

**Notes:** Values are n (%) unless otherwise indicated. A sample-taxon observation is one taxon observed within one sample. Consensus categories indicate the number of the six Discovery runs assigning that observation to its modal reference-aligned context. Context-specific rows include observations with a unique consensus context; ties are shown separately so that rows reconcile with the overall denominator. Pairwise exact agreement was calculated over the 15 unique run pairs using the common-set denominator. Reference run denotes original Trial 1; original Trials 2-6 are reported as Trials 1-5.

### Supplementary Table 4. Distribution of retained ordinal samples across original study timepoints

| Ordinal sample | Timepoint 1 | Timepoint 2 | Timepoint 3 | Timepoint 4 | Timepoint 5 | Timepoint 6 |
|---|---|---|---|---|---|---|
| Visit 1 | 95 (90.5%) | 9 (8.6%) | 0 (0.0%) | 1 (1.0%) | 0 (0.0%) | 0 (0.0%) |
| Visit 2 | 0 (0.0%) | 73 (69.5%) | 16 (15.2%) | 11 (10.5%) | 1 (1.0%) | 4 (3.8%) |

Values are n (% within ordinal sample) among the 105 participants contributing both retained samples. Ordinal sample denotes the first or second observed stool sample after within-participant ordering by collection day and does not necessarily correspond to a fixed scheduled study timepoint. Collection day is reported as median (range).

### Supplementary Table 5. Performance of matched microbiome representations for recurrence classification

| Metric | CLR abundance | Thresholded presence | Taxon-in-context |
|---|---|---|---|
| AUROC | 0.543 (0.424 to 0.663) | 0.536 (0.415 to 0.661) | 0.707 (0.600 to 0.805) |

| | | | |
|---|---|---|---|
| Average precision (PR-AUC) | 0.413 (0.312 to 0.555) | 0.460 (0.338 to 0.618) | 0.644 (0.503 to 0.780) |
| Brier score | 0.274 (0.239 to 0.311) | 0.278 (0.238 to 0.318) | 0.221 (0.198 to 0.246) |
| Log loss | 0.760 (0.678 to 0.845) | 0.772 (0.679 to 0.868) | 0.634 (0.585 to 0.689) |
| Sensitivity | 0.524 (0.368 to 0.676) | 0.571 (0.410 to 0.727) | 0.714 (0.571 to 0.853) |
| Specificity | 0.635 (0.508 to 0.754) | 0.571 (0.448 to 0.690) | 0.508 (0.387 to 0.632) |
| PPV | 0.489 (0.333 to 0.634) | 0.471 (0.327 to 0.609) | 0.492 (0.367 to 0.617) |
| NPV | 0.667 (0.542 to 0.787) | 0.667 (0.538 to 0.796) | 0.727 (0.595 to 0.861) |
| Balanced accuracy | 0.579 (0.479 to 0.681) | 0.571 (0.470 to 0.671) | 0.611 (0.520 to 0.704) |

All models used the same 105 participants (42 with recurrence and 63 without recurrence; recurrence prevalence, 40%), 223 taxa and two ordinal samples per participant. Values are point estimates with conditional 95% confidence intervals in parentheses, rounded to three decimal places. Intervals were estimated by percentile bootstrap using 2,000 common participant-level resamples of pooled outer-fold predictions, without refitting the prediction pipelines. Recurrence was the positive class. Sensitivity, specificity, PPV, NPV and balanced accuracy used outer-test classifications at fold-specific thresholds selected from inner out-of-fold training predictions by maximizing balanced accuracy and locked before outer-test evaluation. PPV and NPV reflect the recurrence prevalence in this cohort. Presence was defined as relative abundance ≥0.1%. AUROC, area under the receiver operating characteristic curve; CLR, centred log ratio; PR-AUC, area under the precision–recall curve, calculated here as average precision; PPV, positive predictive value; NPV, negative predictive value. Lower Brier score and log loss indicate better probability accuracy.

### **Supplementary Table 6. Recurrence discrimination across five linear classifiers**

| **Classifier** | **Context AUROC** | **CLR AUROC** | **Presence AUROC** | **Context − CLR ΔAUROC (95% CI)** | **Context − presence ΔAUROC (95% CI)** |
|---|---|---|---|---|---|

| | | | | | |
|---|---|---|---|---|---|
| L2 logistic regression | 0.653 | 0.465 | 0.543 | 0.188 (0.077 to 0.304) | 0.110 (0.015 to 0.211) |
| L1 logistic regression | 0.685 | 0.454 | 0.520 | 0.231 (0.091 to 0.375) | 0.164 (0.036 to 0.295) |
| Elastic-net logistic regression | 0.706 | 0.483 | 0.568 | 0.223 (0.100 to 0.352) | 0.138 (0.033 to 0.249) |
| Linear support vector machine | 0.683 | 0.487 | 0.570 | 0.196 (0.072 to 0.321) | 0.113 (0.013 to 0.213) |
| Shrinkage linear discriminant analysis | 0.665 | 0.486 | 0.539 | 0.179 (0.061 to 0.301) | 0.126 (0.027 to 0.229) |

All comparisons used the same 105 participants (42 with recurrence), two ordinal samples and a common 226-taxon universe. AUROCs were calculated from participant-level out-of-fold scores averaged across 20 repetitions of nested five-fold cross-validation with three-fold inner validation. ΔAUROC denotes context AUROC minus comparator AUROC. Paired conditional 95% confidence intervals were estimated using 2,000 common participant-level percentile bootstrap resamples of the averaged scores, without model refitting or adjustment for multiple comparisons. Linear support vector machine scores were sigmoid-transformed margins without fitted probability calibration. Presence was defined as relative abundance ≥0.1%. The supplementary CLR calculation used the reclosed 226-taxon subcomposition and differs from the primary analysis in Table 5. AUROC, area under the receiver operating characteristic curve; CI, confidence interval; CLR, centred log ratio.

### **Supplementary Table 7. Stable taxon-in-context predictors in the context-only recurrence model**

| **Taxon** | **Context** | **Visit** | **Selected fits (%)** | **Positive/ negative fits** | **Sign consistency (%)** | **Median β** | **Group** |
|---|---|---|---|---|---|---|---|
| *Eubacterium eligens* | BV | 1 | 5/5 (100%) | 5/0 | 100% | 0.229 | R |
| *Bacteroides xylanisolvens* | BV | 2 | 5/5 (100%) | 0/5 | 100% | −0.136 | N |
| *Parabacteroides goldsteinii* | BV | 2 | 5/5 (100%) | 5/0 | 100% | 0.146 | R |
| *Streptococcus parasanguinis* | PC | 2 | 5/5 (100%) | 5/0 | 100% | 0.178 | R |
| *Bacteroides xylanisolvens* | BV | 1 | 4/5 (80%) | 0/4 | 100% | −0.111 | N |
| *Clostridium innocuum* | BV | 1 | 4/5 (80%) | 0/4 | 100% | −0.171 | N |

| | | | | | | | |
|---|---|---|---|---|---|---|---|
| Ruminococcaceae bacterium D5 | PC | 1 | 4/5 (80%) | 4/0 | 100% | 0.118 | R |
| Firmicutes bacterium CAG 110 | PC | 1 | 4/5 (80%) | 4/0 | 100% | 0.119 | R |
| Ruminococcaceae bacterium D5 | PC | 2 | 4/5 (80%) | 4/0 | 100% | 0.118 | R |
| *Alistipes finegoldii* | BV | 2 | 3/5 (60%) | 0/3 | 100% | −0.158 | N |
| *Hungatella hathewayi* | BV | 2 | 3/5 (60%) | 0/3 | 100% | −0.185 | N |
| *Lactobacillus fermentum* | BL | 1 | 3/5 (60%) | 3/0 | 100% | 0.202 | R |
| *Alistipes indistinctus* | PC | 1 | 3/5 (60%) | 3/0 | 100% | 0.150 | R |
| *Methanobrevibacter smithii* | PC | 2 | 3/5 (60%) | 3/0 | 100% | 0.048 | R |
| *Streptococcus salivarius* | PC | 2 | 3/5 (60%) | 3/0 | 100% | 0.164 | R |

Stable features were selected in ≥60% of the five outer fits of the primary context-only model and showed ≥80% coefficient-sign consistency among nonzero coefficients. Selected fits are reported as the number of fits selecting the feature out of five, with selection percentage in parentheses. Positive/negative fits count selected fits with positive or negative coefficients. Sign consistency is the percentage of nonzero coefficients with the majority sign. β denotes the median standardized logistic-regression coefficient across selected fits. Group indicates association with recurrence (R) or non-recurrence (N); positive and negative coefficients correspond to R and N, respectively. Context codes: BV, Mixed *B. vulgatus*; PC, Complementary *P. copri*; BL, Concentrated *B. longum*. These coefficients describe associations rather than causal effects.

### **Supplementary Table 8. Fit-to-fit reproducibility of microbial-context architectures across candidate K**

| **Number of contexts, K** | **Fits per configuration** | **Within-configuration fit pairs, n** | **Matched architecture distance, median (IQR)** | **90th percentile** |
|---|---|---|---|---|
| 2 | 20 | 4,940 | 0.000 (0.000-0.541) | 0.544 |
| 3 | 20 | 4,940 | 0.027 (0.003-0.130) | 0.296 |
| 4 | 20 | 4,940 | 0.026 (0.009-0.037) | 0.166 |

| | | | | |
|---|---|---|---|---|
| 5 | 15 | 2,730 | 0.086 (0.061-0.117) | 0.151 |
| 6 | 10 | 1,170 | 0.106 (0.083-0.133) | 0.156 |
| 7 | 8 | 728 | 0.112 (0.088-0.133) | 0.155 |
| 8 | 6 | 390 | 0.093 (0.077-0.112) | 0.129 |
| 9 | 5 | 260 | 0.083 (0.062-0.104) | 0.123 |
| 10 | 4 | 156 | 0.066 (0.054-0.083) | 0.093 |

**Note.** Fit-to-fit architecture distance was calculated only between Discovery fits obtained under identical K, abundance-threshold and prevalence-threshold settings. For each fit pair, contexts were optimally matched to minimize the mean Euclidean distance between their two-dimensional PCoA coordinates. Values are median (interquartile range; IQR) across all within-configuration fit pairs; the 90th percentile is shown as an additional measure of the upper tail. Lower distances indicate greater architectural reproducibility. The number of sampled fits per configuration decreased for $K > 4$.

### Supplementary Data 1. Age-associated context-responsive taxa and context pivots
